# Phylogenetic community structure metrics and null models: a review with new methods and software

**DOI:** 10.1101/025726

**Authors:** Eliot T. Miller, Damien R. Farine, Christopher H. Trisos

## Abstract

Competitive exclusion and habitat filtering are believed to have an important influence on the assembly of ecological communities, but ecologists and evolutionary biologists have not reached a consensus on how to quantify patterns that would reveal the action of these processes. No fewer than 22 phylogenetic community structure metrics and nine null models can be combined, providing 198 approaches to test for such patterns. Choosing statistically appropriate approaches is currently a daunting task. First, given random community assembly, we assessed similarities among metrics and among null models in their behavior across communities varying in species richness. Second, we developed spatially explicit, individual-based simulations where communities were assembled either at random, by competitive exclusion or by habitat filtering. Third, we quantified the performance (type I and II error rates) of all 198 approaches against each of the three assembly processes. Many metrics and null models are functionally equivalent, more than halving the number of unique approaches. Moreover, an even smaller subset of metric and null model combinations is suitable for testing community assembly patterns. Metrics like mean pairwise phylogenetic distance and phylogenetic diversity were better able to detect simulated community assembly patterns than metrics like phylogenetic abundance evenness. A null model that simulates regional dispersal pressure on the community of interest outperformed all others. We introduce a flexible new R package, *metricTester*, to facilitate robust analyses of method performance. The package is programmed in parallel to readily accommodate integration of new row-wise matrix calculations (metrics) and matrix-wise randomizations (null models) to generate expectations and quantify error rates of proposed methods.

## Introduction

“…we may be studying an attribute about which we cannot be sure what measurements can actually represent it or even whether a hypothesized attribute actually exists” (Houle et al. 2011).

The idea that competition among species increases with relatedness goes back at least to Darwin (1859), who noted that more closely related species tend to be more ecologically similar and should therefore compete more intensely (reviewed in Cavender-Bares et al. 2009). Referred to as the competition-relatedness hypothesis (Cahill et al. 2008), competitive exclusion is predicted to result in the co-occurrence of less closely related species than would be expected if communities were assembled entirely via stochastic processes (phylogenetic overdispersion; Elton 1946, Webb et al. 2002, but see Mayfield and Levine 2010), such as speciation and dispersal. In contrast to competitive exclusion, habitat filtering is the process whereby only those species possessing similar traits are able to survive and reproduce within a given abiotic environment (Harper 1977, Keddy 1992). Thus, to the extent that such traits are evolutionarily conserved, habitat filtering results in local assemblages of species more closely related than expected by chance (phylogenetic clustering; Webb 2000, Cavender-Bares et al. 2009). Habitat filtering operates largely independently of individual interactions, whereas competitive exclusion occurs via either direct or indirect agonistic interactions among individuals of different species. Until the 1990s, few methods existed to test for patterns of relatedness within communities, and those available took a taxonomic rather than a phylogenetic approach (Elton 1946, Vane-Wright et al. 1991).

Over the past 25 years a large number of metrics have been developed to quantify phylogenetic patterns in community structure, by which one might infer the action of community assembly processes. However, misconceptions about the relationships of these metrics to each other and to species richness (reviewed in Box 1) have reduced their impact on our understanding of community assembly. The link between our theories of community assembly and our ostensible measures of it are tenuous, and the measures themselves are not well understood (Houle et al. 2011). Furthermore, while the metrics introduced by Webb (Webb 2000, Webb et al. 2002) have been most influential in community ecology, many other metrics have also received widespread use, and yet their mathematical properties and performance across different community assembly processes has not been comprehensively assessed. Recent reviews (Kraft et al. 2007, Kembel 2009, Vamosi et al. 2009, Vellend et al. 2011, Pearse et al. 2014) have addressed metric performance, but have evaluated only partially overlapping sets of metrics, often using different methods and classification schemes. Consequently, results cannot easily be compared among studies, making the selection of appropriate metrics for empirical research difficult.

### Box 1

#### Abbreviated history of phylogenetic community structure metrics

Faith (1992) introduced PD, a metric that quantifies the unique evolutionary history represented by co-occurring taxa. It was intended (and is often used) as a conservation tool. While PD built upon previous work by Vane-Wright et al. (1991) and others, it was the first to explicitly incorporate phylogeny. Since PD is the sum of all branch lengths connecting the species in a community (Table 1), the assumption that it increases with additional species, and is therefore correlated with species richness, was implicit (exact solution provided by Nipperess & Matsen 2013).

Subsequently, Clarke and Warwick introduced metrics (Δ, Δ+, Δ*) focused on the average branch length among a group of taxa or individuals, again linking their methodology to conservation decisions (Warwick and Clarke 1995, 1998, Clarke and Warwick 1998, 1999). Their pioneering papers explored some statistical properties of the metrics, including the fact that mean expected Δ+ is not correlated with species richness, but the width of its confidence intervals decreases with increasing species richness (creating a “confidence funnel”). Yet, the conservation-specific scope of their papers limited their impact on community ecology.

Webb (2000) introduced two new metrics--MPD and MNTD--and the standardized forms of these, NRI (net relatedness index) and NTI (nearest taxon index). Initially, MPD was slightly different than Clarke and Warwick’s metrics, only incorporating nodal distances, but by Webb et al. (2002) the definition had expanded to incorporate branch length, and was therefore equivalent to Δ+ (Appendix S2). Yet, by linking community assembly processes with these phylogenetic patterns, it was MPD and MNTD that revolutionized the field of community ecology. Moreover, despite the equivalency of MPD and Δ+, Webb stated that both MPD and MNTD are correlated with species richness when only MNTD is (Fig. 1A), and devised standardization procedures to “correct” for this. This misperception occasionally persists to the present (e.g., Ulrich & Fattorini 2013), despite empirical solutions to the contrary (Tsirogiannis and Sandel 2013).

Helmus et al. (2007) introduced PSE, the “first” metric to incorporate abundance information. While this is not entirely true (Rao 1982, Warwick and Clarke 1995, Hardy and Senterre 2007), their focus on community assembly linked PSE with quintessential evolutionary questions. Helmus et al. (2007) also introduced two other metrics intended to be similar but superior to NRI and NTI--PSV and PSC. The noted advantage to these is the lack of need for a reference species pool, and therefore the ability of these metrics to transcend the particulars of the phylogeny and community data matrix at hand, and allow raw metric values to be directly compared. However, these should therefore have been compared with MPD and MNTD, respectively. Had this been done, it would have been noted that PSV and PSC are directly proportional to MPD and MNTD, respectively, a still all but unknown fact (though see Vellend et al. 2011). Instead, PSV and PSC were compared with NRI and NTI. As a further complication, the PSC function in *picante* (Kembel et al. 2010) returns the inverse of PSC (M. Helmus, pers. comm.). This, and a belief of uncertain affinities that PSC is not inherently correlated with species richness, confounded subsequent papers (e.g. Giehl & Jarenkow 2012; Villalobos et al. 2013).

Cadotte et al. (2010) introduced metrics focused on phylogenetic abundance distributions. We review seven of those here: PD_*c*_ (discussed earlier, Faith (2007)), PAE, IAC, ED, *H*_ED_, *E*_ED_, *H*_AED_, and *E*_AED_ (see Table 1). Cadotte et al. (2010) showed their metrics ranked communities differently than each other and than metrics like PSV and MNTD, but offered no discussion of the metrics’ statistical properties, nor has any subsequent paper. The metrics are available in *ecoPD* (http://r-forge.r-project.org/projects/ecopd/).

*We discuss six additional metrics in this paper: QE (Rao 1982), SimpsonsPhy (Hardy and Senterre 2007), abundance-weighted (AW) MNTD, and three variants of AW MPD (Table 1, Appendix S2). Both complete AW MPD and AW MNTD were introduced in Phylocom* (Webb et al. 2008) and *picante* without accompanying publication, and their statistical properties and relationship to other metrics remains essentially unknown. Interspecific AW MPD was introduced in (Miller et al. 2013), and intraspecific AW MPD is “first” described in the current paper (Appendix 2), though as we subsequently discovered, it is equivalent to Δ (Clarke and Warwick 1998). Similarly, after exploring the behavior of QE and SimspsonsPhy and finding them equivalent, we realized this was already known (Hardy and Senterre 2007, Allen et al. 2009).

Statistically evaluating the significance of an observed phylogenetic community structure metric requires a null expectation. Thus, since their introduction, phylogenetic community metrics have been linked to null models (Webb 2000), when in fact, they are independent concepts. This conceptual link has led to the creation of redundant metrics and frequent and continuing confusion in the literature (Box 1). We suggest that practitioners should consider phylogenetic community structure methods as a set of possible metrics (e.g., row-wise calculations) and a set of possible null models (e.g., repeated matrix-wise randomizations), any of which can be combined to create a unique metric + null model approach. Thus, the metric value for a particular community and phylogeny is fixed, but the significance of that metric varies according to which null model is used (Connor and Simberloff 1979, Diamond and Gilpin 1982, Gotelli 2000). A good null model randomizes those structures in the observed data (e.g., individual co-occurrence patterns) relevant to the null hypothesis, and maintains structures in the dataset unrelated to the null hypothesis (e.g., species’ abundance distributions) (Gotelli and Graves 1996). In practice, null model performance, specifically type I (false positive) and II (false negative) error rates, and redundancy among null models is rarely tested (but see e.g., Gotelli 2000, Kembel 2009).

Here, we compare the performance of 22 phylogenetic community structure metrics (Table 1) and 9 null models (Table 2). We develop spatially explicit, individual-based simulations of community assembly due to habitat filtering, competitive exclusion or the random placement of individuals, and then compare the ability (type I and II error rates) of each metric + null model combination to identify the correct assembly process. We document cases of equivalency among metrics and null models. We also assess the response of both metrics and null models to variation in species richness. We conclude by discussing the implications of our findings for future tests of community assembly processes.

**Table 1.** The 22 phylogenetic community structure metrics reviewed in this paper. We paraphrase (or sometimes directly quote) the original description of the metric. While some metrics we discuss are in fact equivalent, these original descriptions often emphasized their uniqueness. IAC is a node-based metric. We multiplied it by -1 such that decreases in its value corresponded with increased clustering.

| Metric | Abbreviation | Description | Citation |
| --- | --- | --- | --- |
| Quadratic entropy | QE | Within community diversity based on species dissimilarity. | (Rao 1982) |
| Phylogenetic diversity | PD | Sum of total branch lengths for a set of species, and length to root if set does not span it. | (Faith 1992) |
| Non-abundance-weighted mean pairwise phylogenetic distance | MPD | Mean of all pairwise branch lengths for a set of species. | (Webb 2000, Webb et al. 2002) |
| Non-abundance-weighted mean nearest taxon distance | MNTD | Mean of the branch lengths separating each species from its closest relative in the set of species. | (Webb 2000, Webb et al. 2002) |
| Taxonomic diversity* | $\Delta$ | Average phylogenetic distance between any two individuals from a set. | (Clarke and Warwick 1998) |
| Taxonomic distinctness* | $\Delta^*$ | Average phylogenetic distance between any two heterospecific individuals. | (Clarke and Warwick 1998) |
| Presence-absence case of taxonomic diversity* | $\Delta^+$ | Average phylogenetic distance between any two species from a set. | (Clarke and Warwick 1998) |
| Phylogenetic species variability | PSV | Measures how phylogenetic relatedness decreases the variance of a hypothetical Brownian motion trait shared by all species in the | (Helmus et al. 2007) |

Table 1 *continued*
| Metric | Abbreviation | Description | Citation |
| --- | --- | --- | --- |
| Phylogenetic species clustering | PSC | Modified form of PSV incorporating maximum off-diagonal element matrix of community phylogenetic correlation structure. | (Helmus et al. 2007) |
| Phylogenetic species evenness | PSE | Modified form of PSV incorporating species abundance. | (Helmus et al. 2007) |
| Phylogenetic form of Simpson's index | SimpsonsPhy | Extension of Simpson diversity index that incorporates phylogenetic information. | (Simpson 1949, Hardy and Senterre 2007) |
| Abundance-weighted MNTD | AW MNTD | Abundance-weighted form of MNTD. | (Webb et al. 2008) |
| Phylogenetic diversity without regard to a larger regional pool | PD<sub>c</sub> | Sum of total branch lengths for a set of species, not including length to root. | (Faith 2007, Cadotte et al. 2010) |
| Phylogenetic abundance evenness | PAE | "Phylogenetic evenness of abundance distribution scaled by branch length." | (Cadotte et al. 2010) |
| Imbalance of abundance | IAC | IAC. "Relative per-node imbalance in individual distribution." | (Cadotte et al. 2010) |
| Community evolutionary distinctiveness | H<sub>ED</sub> | "Entropic measure of diversity of evolutionary distinctiveness among species." | (Cadotte et al. 2010) |
| Equitability evolutionary distinctiveness | E<sub>ED</sub> | "Equitability of H<sub>ED</sub>." | (Cadotte et al. 2010) |
| Community abundance-weighted evolutionary | H<sub>AED</sub> | "Entropic measure of diversity of evolutionary distinctiveness among individuals." | (Cadotte et al. 2010) |

Table 1 *continued*
| Metric | Abbreviation | Description | Citation |
| --- | --- | --- | --- |
| Equitability<br>abundance-weighted<br>evolutionary<br>distinctiveness | $E_{\text{AED}}$ | “Equitability of $H_{\text{AED}}$.” | (Cadotte et al. 2010) |
| Complete<br>abundance-weighted<br>MPD | complete AW<br>MPD | An abundance-weighted form of MPD. Average phylogenetic distance between two individuals from a set, possibly between the same individual. | (Webb et al. 2008, Appendix S2 of this paper) |
| Intraspecific<br>abundance-weighted<br>MPD | intra AW<br>MPD | An abundance-weighted form of MPD. Average phylogenetic distance between any two individuals from a set. | (Appendix S2 of this paper) |
| Interspecific<br>abundance-weighted<br>MPD | inter AW<br>MPD | An abundance-weighted form of MPD. Average phylogenetic distance between two heterospecific individuals. | (Miller et al. 2013, Appendix S2 of this paper) |

\* Denotes three metrics not directly assessed here due to equivalency with other metrics
(see Appendix S2), leaving 19 focal metrics in this paper.

**Table 2.**
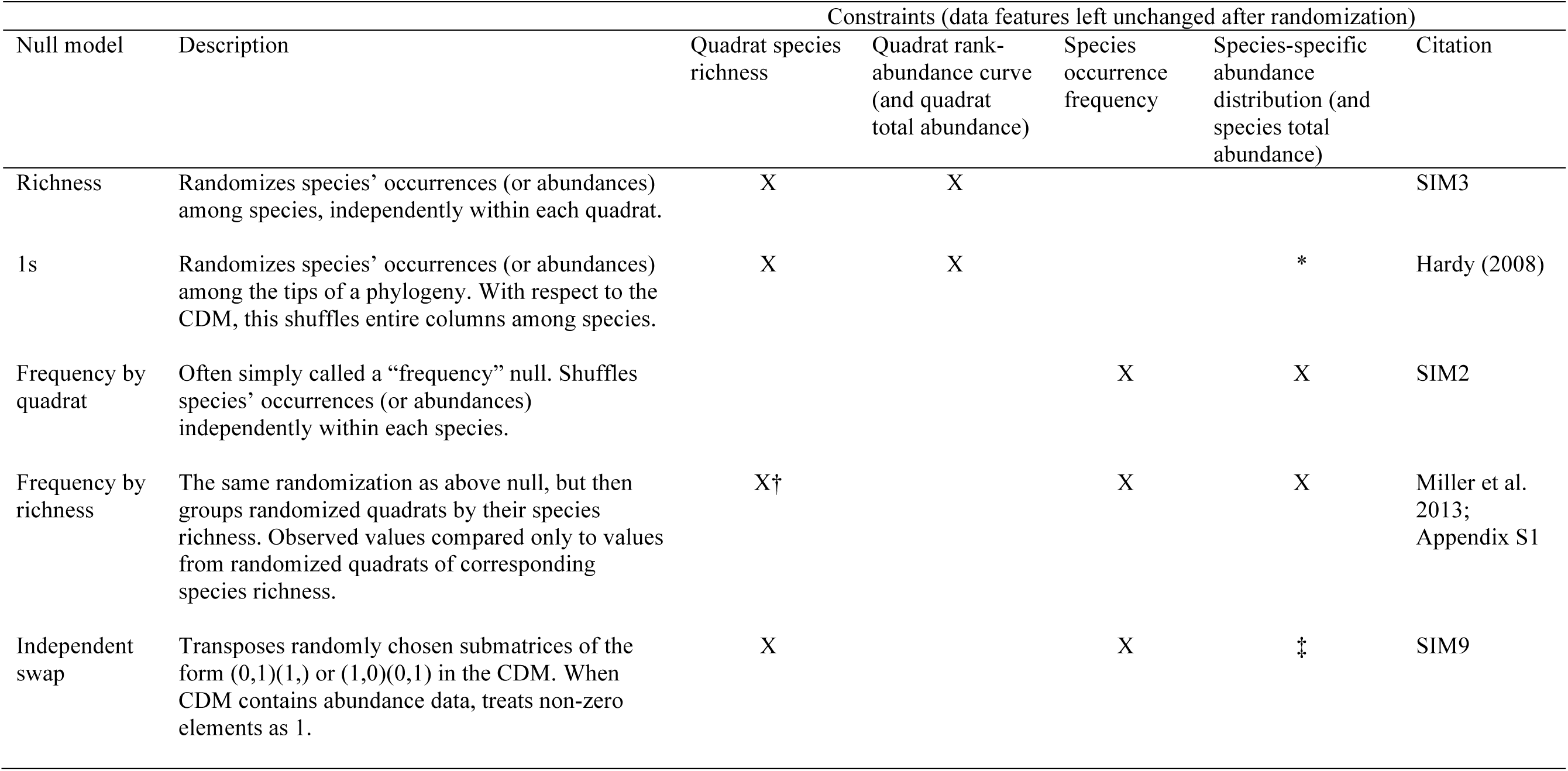
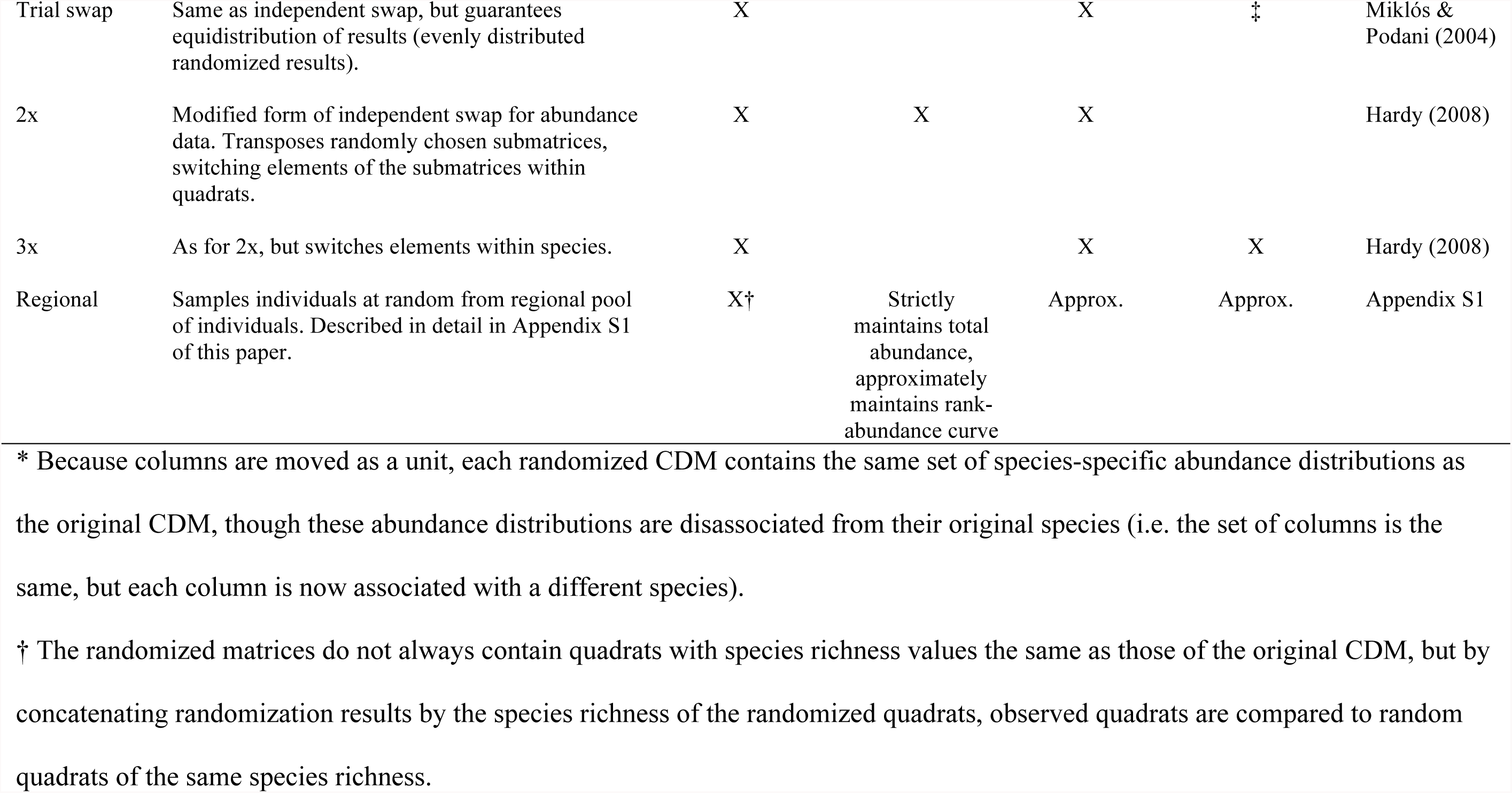
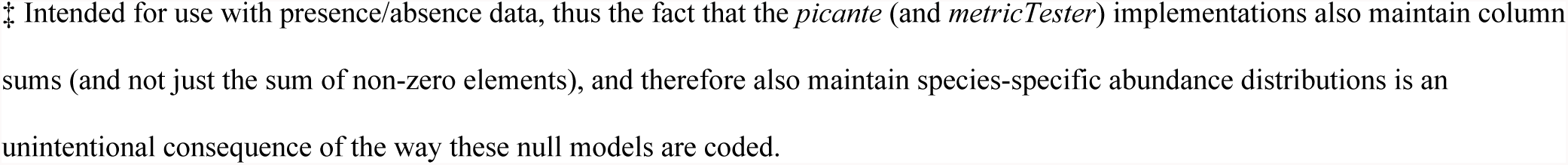
The nine null models reviewed in this paper. A community data matrix (CDM) where quadrats (i.e. sites or samples) are rows and species are columns is used as the input. The citation lists either the simulation name from Gotelli (2000), or gives a more recent citation where necessary.

## Methods

We adopt the following terminology. The community is the spatial extent (i.e. study area) of interest. The quadrat is the sampling unit. For instance, 20, 1-ha forest plots in the Ecuadorian Amazon would be considered 20 quadrats of the rainforest community. We refer to the quadrat (row) by species (column) data matrix as the community data matrix (CDM).

### Null model background

We tested the performance of nine null models (Table 2). While distinctions are often drawn between models that randomize phylogenetic tip labels and those that randomize the CDM (e.g., Hardy 2008), this distinction is false; all tip-shuffling null models can be performed by matrix shuffling (Table 2, Appendix S1).

Perhaps the simplest of the null models we tested is the richness model, which shuffles species occurrences or abundances randomly within quadrats (rows), thereby maintaining species richness (row totals) and, for abundance data, total abundance and the rank-abundance curve of each quadrat.

In contrast, the frequency null model shuffles species occurrences within species (columns) in the CDM, which maintains the occurrence frequency or total abundance of each species (column totals), but not quadrat species richness. Instead, per-quadrat randomized species richness values are distributed around the mean per-quadrat species richness in the observed CDM. Species-poor quadrats will tend to be compared with quadrats of higher species richness, which does not incorporate the large variance expected of randomized metric scores derived from repeated small draws from the species pool (Efron 1979), and thus this model may exhibit elevated error rates. We refer to this null model as the “frequency by quadrat”, to distinguish it from another model described below.

The independent swap null model was developed to reduce error rates by maintaining both species occurrence frequencies and quadrat species richness (row and column totals) (Gotelli 2000, Gotelli and Entsminger 2001). The trial swap (Miklós and Podani 2004) was subsequently introduced as a more efficient approach to maintain the same structures in the null model. We used 10^5^ swaps for these algorithms (Fayle and Manica 2010). In addition, Miller et al. (2013, Appendix 3 of that paper) developed the “frequency by richness” null model which, like the frequency null, shuffles occurrences within species but then concatenates the randomized quadrats by their species richness values, thereby maintaining species occurrence frequencies and quadrat species richness.

Prior to the development of abundance-weighted metrics, few null models intentionally maintained features of abundance distributions. For example, a species might occur infrequently but in large numbers. Hardy (2008) introduced the 2x and 3x null models to maintain both species richness and occurrence frequency, as well as either the species or quadrat-level structure of abundance data. The 2x maintains the total abundance and rank-abundance curve of each quadrat, but neither species’ abundances nor the set of species-specific abundance distributions. In contrast, the 3x maintains species’ abundances and the set of species-specific abundance distributions, but not the abundance distributions of each quadrat. No null model that we know of maintains species richness, species occurrence frequency, species-specific and quadrat-specific abundance distributions (it is likely not possible via matrix shuffling).

We developed (Appendix S1) and tested a model that approximates this behavior, which we call the regional null. The regional null simulates dispersal of individuals into the local community from the regional pool, where local dynamics have no influence on the regional pool. Instead of using observed species abundance and occurrence frequencies from the community (i.e. study area) of interest, individuals are drawn randomly from an abundance-informed regional pool such that species’ colonization probabilities are proportional to regional abundances (Lessard et al. 2012).

### metricTester

We wrote an R software package to run our analyses. *metricTester* is available from GitHub, along with associated documentation, and can be installed using the *devtools* package (<u>https://github.com/eliotmiller/metricTester</u>). To eliminate conflicts with *picante* (Kembel et al. 2010) we renamed some of the functions in *ecoPD* (Cadotte et al. 2010) and rebuilt the package, hosted under the name *ecoPDcorr* in the same Github account. *metricTester* interfaces with additional packages (Paradis et al. 2004, Eastman et al. 2011, Pennell et al. 2014), and is programmed in parallel and designed to facilitate the addition of new metrics, null models and community simulations. Thus, the performance of proposed metrics and null models can be tested against community simulations of the user’s choice. Generation of such expectations is not limited to phylogenetic community structure methods, and extends to any row-or column-wise metric calculation with repeated matrix-wise null model randomization.

### General behavior of the metrics

To understand the behavior of the 19 focal metrics (Table 1) across variation in species richness, we generated a phylogenetic tree that terminated at 50 species using a pure-birth model (birth=0.1), then assembled a CDM that included one “quadrat” at every species richness value between 10 and 40 species (thus, each CDM had 31 rows). Quadrats were created by randomly sampling species from the phylogeny, and then assigning these species abundances from a log-normal distribution (mean = 3, SD = 1). We then calculated and retained focal metrics for each quadrat. Using the same phylogeny, we repeated the process of filling a new CDM, calculating and retaining all metric values 50,000 times. We then calculated the mean and 95% confidence intervals (CI) of each metric at each species richness value, and plotted these across their respective species richness values.

We used Pearson correlations to assess metric similarities and identify redundancies among the metrics. Because of the large number of simulations, some metrics that appear exactly correlated do in fact differ subtly (Appendix S2). We used these correlations to derive a distance matrix, then clustered metrics with a complete linkage method, and used this to generate a dendrogram.

### General behavior of the null models

To identify similarities among null models, we explored the behavior of 9 null models (Table 2) across variation in species richness. We generated CDMs as above, except that species richness values ranged from 10 to 25. We used non-abundance-weighted mean pairwise phylogenetic distance (MPD) for these analyses because it is not inherently correlated with species richness (Fig. 1A), and therefore does not confound metric and null model behavior across species richness. In addition, null model expectations of MPD converged relatively quickly. Using an abundance-weighted metric did not affect results (not shown). To identify null models that do or do not converge efficiently on a stable range of expected metric values, we explored how null model expectations changed with increasing numbers of randomizations of the CDM (Appendix S1). We did this by plotting the expected CI across the corresponding species richness while increasing the randomization of a given, initial CDM and phylogeny.

### Individual-based spatial simulations of community assembly to assess the performance of metric + null combinations

The first two sets of analyses illustrated the general behavior of each metric and null model. In this third analysis, we assessed the ability of each metric + null model combination to detect a given assembly process. Because of the large number of steps in this analysis, we include a schematic to aid the following explanation (Appendix S3).

Total computing time required to run these tests (>7 years) precluded systematic examination of sensitivity to simulation parameters, but results were very similar across preliminary exploration of parameter space (Appendix S4).

To generate test cases against which to assess each metric + null approach, we simulated three types of spatially explicit communities, intended to model random assembly and the extremes of habitat filtering and competitive exclusion. Each spatial simulation produced a 316 × 316 m (10 ha) community, and 1,009 such communities of each type were generated. We began by generating a phylogeny of 100 species using a pure-birth model (birth = 0.1) and log-normal rank abundance curve, and randomly assigned species abundances from this distribution. We expanded assigned abundances to create a vector of individuals with species identities. In the random assembly spatial simulation, these individuals were then randomly placed within the community.

In habitat filtering simulations, we independently evolved two traits according to a Brownian motion evolutionary process (σ^2^ = 0.1). These traits are meant to mimic two independently evolving environmental preferences, e.g., soil moisture and pH. In our case, we treated these as spatial preferences (i.e. × and y-axis preferences), and scaled the simulated traits to match community bounds. We further smoothed species’ spatial preferences, which initially approximated a normal distribution, to a uniform distribution, such that species’ preferences were evenly distributed but phylogenetically conserved across the arena. We then placed individuals near their spatial preference, with a controllable degree of variation (exact parameters in Appendix S4). This simulation has the effect of placing related individuals near each other in space.

In competitive exclusion simulations, we first placed individuals using the random assembly process. Following this, each generation, we calculated the mean relatedness of every individual in the community to all individuals within 20 m, which we term the “interaction distance”. We then identified the 20% of individuals with the highest mean relatedness. For each of these individuals, we identified the individual within their interaction distance to which they were most closely related, and then randomly selected one of the two individuals to be removed from the community. At the end of each generation, the same number of individuals as was removed was drawn from the original vector of individuals, and situated randomly in the community. This was repeated for 60 generations for each competitive exclusion simulation. Preliminary analyses indicated that results were similar across different interaction distances and percentages of individuals considered (Appendix S4). All spatial simulations employed 200-400 individuals/ha, which is somewhat less than stem-density in Australian tropical rain forests (Murphy et al. 2013), and notably less than those in Ecuador (Valencia et al. 2004). Results were nearly identical, however, when we performed the same analysis with comparable numbers of individuals (Appendix S4).

After each spatial simulation, we generated a CDM by situating 20, non-overlapping quadrats of 31.6 × 31.6 m (0.1 ha) at random and recording the individuals in each quadrat. We then calculated observed metrics. Each CDM was then randomized 1,000 times according to each null model, and for each randomization we calculated and retained all metric values. We then calculated standardized effect scores (SES, e.g., standardized MPD equals NRI, Box 1) and 95% CI per quadrat and per unique species richness value. In other words, randomized results were both concatenated by species richness and by quadrat, and SES and CI calculated for each such approach (Appendix S3). Because results were similar, we generally report results for the quadrat concatenation in the main text, with results from the richness method in Appendix S6. As this is the distinguishing feature between the frequency by richness and frequency by quadrat null models, however, we report these separately in the main text. The regional null is designed to be concatenated by richness, so quadrat method results for this model were discarded.

We used Wilcoxon signed-rank tests to assess error rates. For a given metric + null model approach, a type I error was recorded if the distribution of SES from a random spatial simulation differed significantly from zero (two-sided test). A type II error was recorded if the distribution of SES from either a filtering or competition simulation was not significantly less or more than zero, respectively (one-sided test). Thus, the overall type I error rate for a given approach is the proportion of the 1,009 random spatial simulations where the set of SES differed from zero. The overall type II error rate is the mean proportion of the filtering and competition simulations where the SES did not differ as expected from zero. We used one minus the type I and II error rates as a measure of overall approach performance. The use of confidence intervals to quantify error rates is discussed in Appendix S6.

## Results

### General behavior of the metrics

We evaluated the behavior of 19 focal community phylogenetic metrics (Table 1) across variation in species richness. MPD, interspecific AW MPD, PSV and PAE were not correlated with species richness (Fig. 1A). Intraspecific AW MPD, complete AW MPD, PSE, IAC, *H*_AED_, *H*_ED_, SimpsonsPhy, PD, PD_*c*_, and QE were positively correlated with species richness. MNTD, AW MNTD, PSC, and *E*_ED_ were negatively correlated with species richness. The intercorrelations among metrics (Fig. 1B, Appendix S5) revealed that: (1) MPD is equivalent to PSV; (2) complete AW MPD is equivalent to SimpsonsPhy and QE, and approximately equal to intraspecific AW MPD (Appendix S2) and to PSE; and (3) PSC is equivalent to MNTD. Moreover, MPD, interspecific AW MPD, and intraspecific AW MPD are equivalent to Δ+, Δ*, and Δ, respectively, of Clarke & Warwick (1998) (Box 1, Appendix S2). Thus, of the 22 metrics in Table 1, only 15 are unique and, if very closely correlated metrics are considered equivalent, only 12 of the metrics are truly unique.

We classified metrics into the following groups based on intercorrelations among them (Fig. 1B): Group 1 metrics quantify the mean relatedness among species; Group 2 metrics focus on the relationship between “evolutionary distinctiveness and abundance” (Cadotte et al. 2010); Group 3 metrics focus on patterns of phylogenetic relatedness among nearest relatives; and Group 4 metrics quantify the total relatedness in an assemblage, and are most closely correlated with species richness.

**Figure 1(next page). (A).**
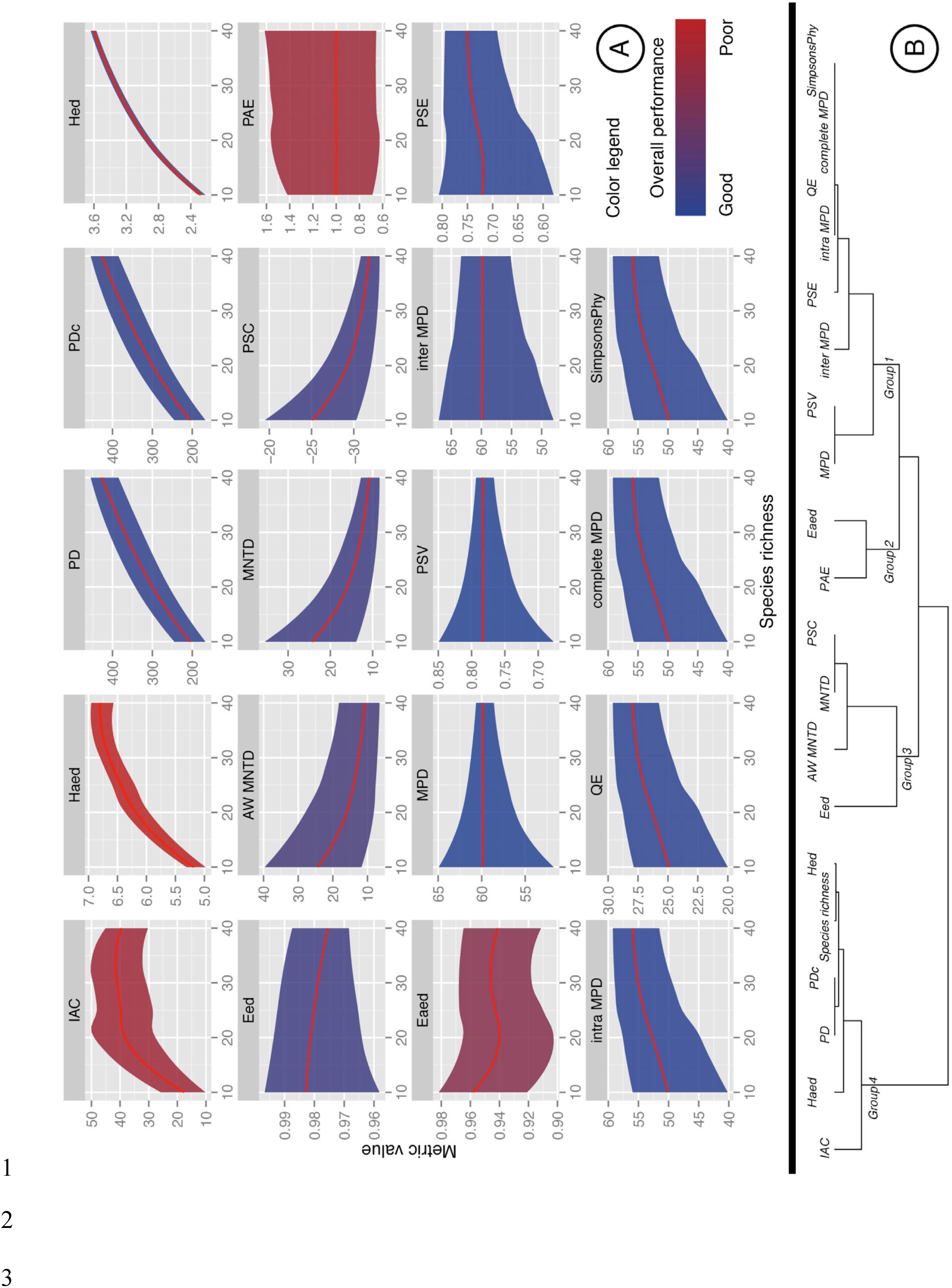
Behavior of 19 focal phylogenetic community structure metrics (Table 1) across variation in species richness. Panels are color-coded from blue (good) to red (poor) according to the mean of type I and II errors across all simulated assembly processes. **(B)** Dendrogram of intercorrelations among the phylogenetic community structure metrics (and species richness itself). Closely correlated metrics are annotated along branches. Group 1 metrics focus on “mean relatedness;” Group 2 metrics on the relationship between “evolutionary distinctiveness and abundance”; Group 3 on “nearest-relative” measures of community relatedness; and Group 4 on “total community diversity” and are particularly closely correlated with species richness.

### General behavior of the null models

The CIs from the richness, 1s, independent swap, trial swap, frequency by richness, and regional null models exhibited confidence funnels (Clarke and Warwick 1998), with more variance observed in smaller (less species rich) samples of the regional species pool (Fig. 2; Fig S1.7). In contrast, the CI of the frequency by quadrat null model did not account for the anticipated increased variance in null model expectations at low species richness, and the value beyond which an observed metric needed to deviate to be considered significant was approximately the same for all quadrats, irrespective of underlying species richness of the quadrat (Fig. 2).

**Figure 2.**
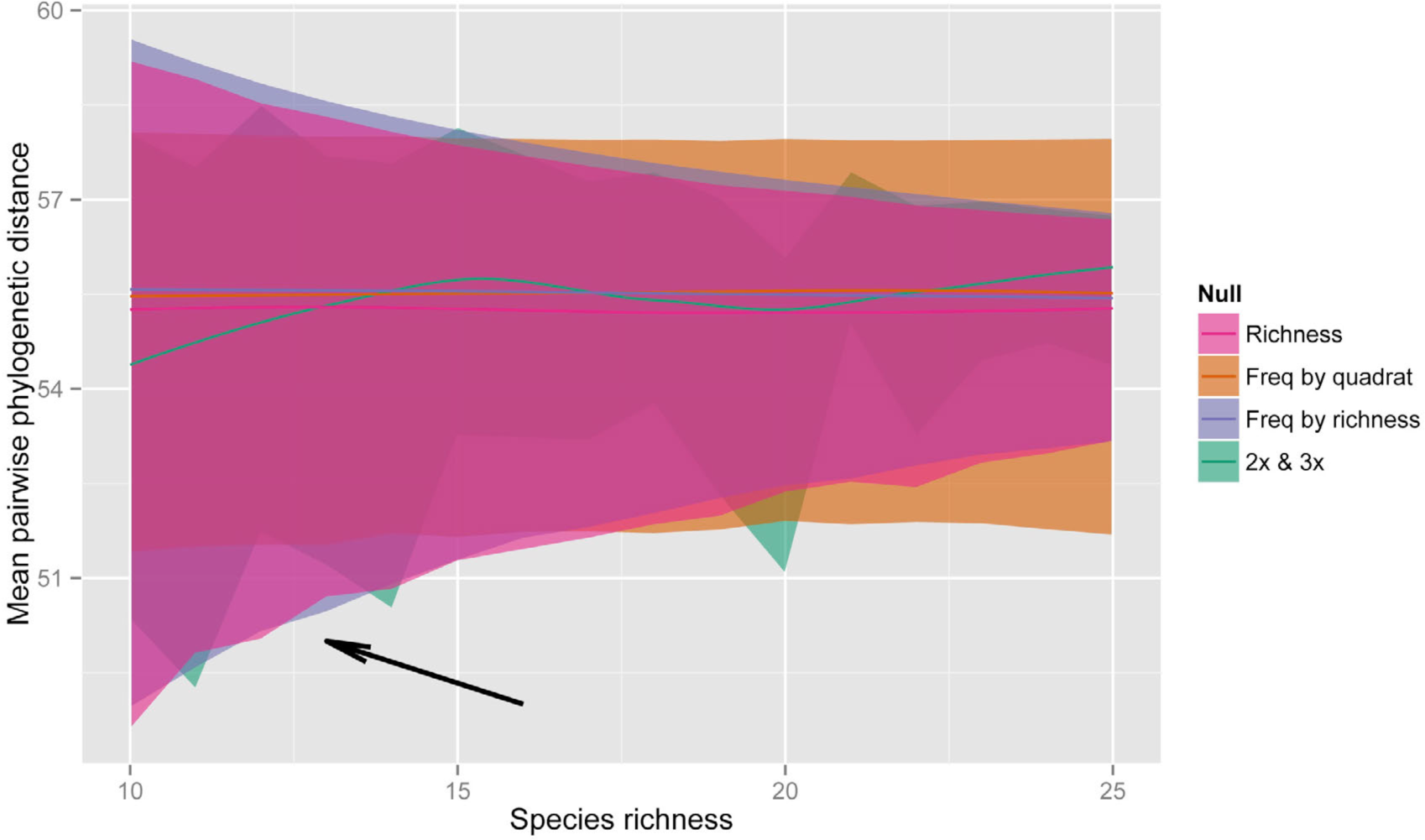
Confidence intervals (95%) for the richness, both forms of the frequency, 2x and 3x null models (Table 2) across variation in species richness. Expectations shown here are the result of 10^5^ randomizations. Because the 2x and 3x nulls follow identical distributions (Fig. S1.5), only a single layer is included in this figure. The arrow indicates a region of particular concern for type I error when using the frequency by quadrat null. Other null model behavior (including the independent swap, trial swap, and regional models) is summarized in Appendix S1.

The 1s and the richness null models converged on very similar null model expectations (Fig. S1.1), showing that despite differences in randomization schemes these models maintain the same elements of the CDM, and they are thus functionally equivalent. This was also the case for the independent swap and the frequency by richness null models (Fig. S1.4). In addition, the trial swap null converged, albeit slowly (after >10^6^ randomizations), on the same expectations as the frequency by richness and independent swap null models (Fig. S1.2). We also found that the expectations when concatenated by richness from the 2x and 3x null models were equivalent, but did not form a confidence funnel (Fig. 2, Fig. S1.5, Appendix S6). Finally, expectations for the independent swap varied depending upon the relationship between species’ occurrence frequency and phylogenetic uniqueness. For instance, when phylogenetically unique species occurred more frequently in the CDM, CI were shifted upwards from models that do not maintain species occurrence frequency (e.g., the richness model, Fig. S1.6).

### Performance of metric + null approaches

We ran 1,009 complete tests (all spatial simulations, null models and metrics). There was a great deal of variation in performance of different metric + null approaches. Across all metrics, for both competitive exclusion and habitat filtering assembly simulations, the frequency by quadrat null showed high rates of type II error, particularly for metrics that were correlated with species richness (Fig. 1A). The 2x and 3x nulls showed high type II error rates, particularly for the detection of habitat filtering and for metrics tailored to be sensitive to differences in abundance distributions (PAE, IAC, *H*_AED_, Fig. 3). The independent swap, trial swap and frequency by richness null models performed reasonably well in habitat filtering simulations when used with some metrics (PD, MPD, MNTD, Fig. 3), but poorly in competitive exclusion simulations with most metrics (Fig. 4). Finally, the richness, 1s and regional nulls performed well with most metrics in both the habitat filtering and competitive exclusion simulations, but the richness and 1s exhibited high type I error rates (Fig. 5).

Focusing on the metrics, PD, PD_*c*_, MNTD and AW MNTD had the greatest power to detect habitat filtering, though Group 1 metrics also performed well (Fig. 3). PD and PD_*c*_ were also relatively powerful at detecting the signature of competitive exclusion (Fig. 4), though here they were outperformed by Group 1 metrics. Group 3 metrics exhibited relatively less power to detect phylogenetic overdispersion, particularly with some null models (3x, independent swap). If we take overall metric performance as the mean of the type I error rates across all null models for the random simulations, and the type II error rates across all null models for the habitat filtering and competitive exclusion simulations, then Group 1 metrics performed best overall, followed closely by PD and PD_c_, and then by Group 3 metrics (Fig. 5). Some metrics (*E*_AED_, PAE, IAC, *H*_AED_) exhibited type I error rates similar to those of the more successful metrics (i.e. 10-11%), but also failed more often than they succeeded to detect simulated community assembly processes.

**Figure 3.**
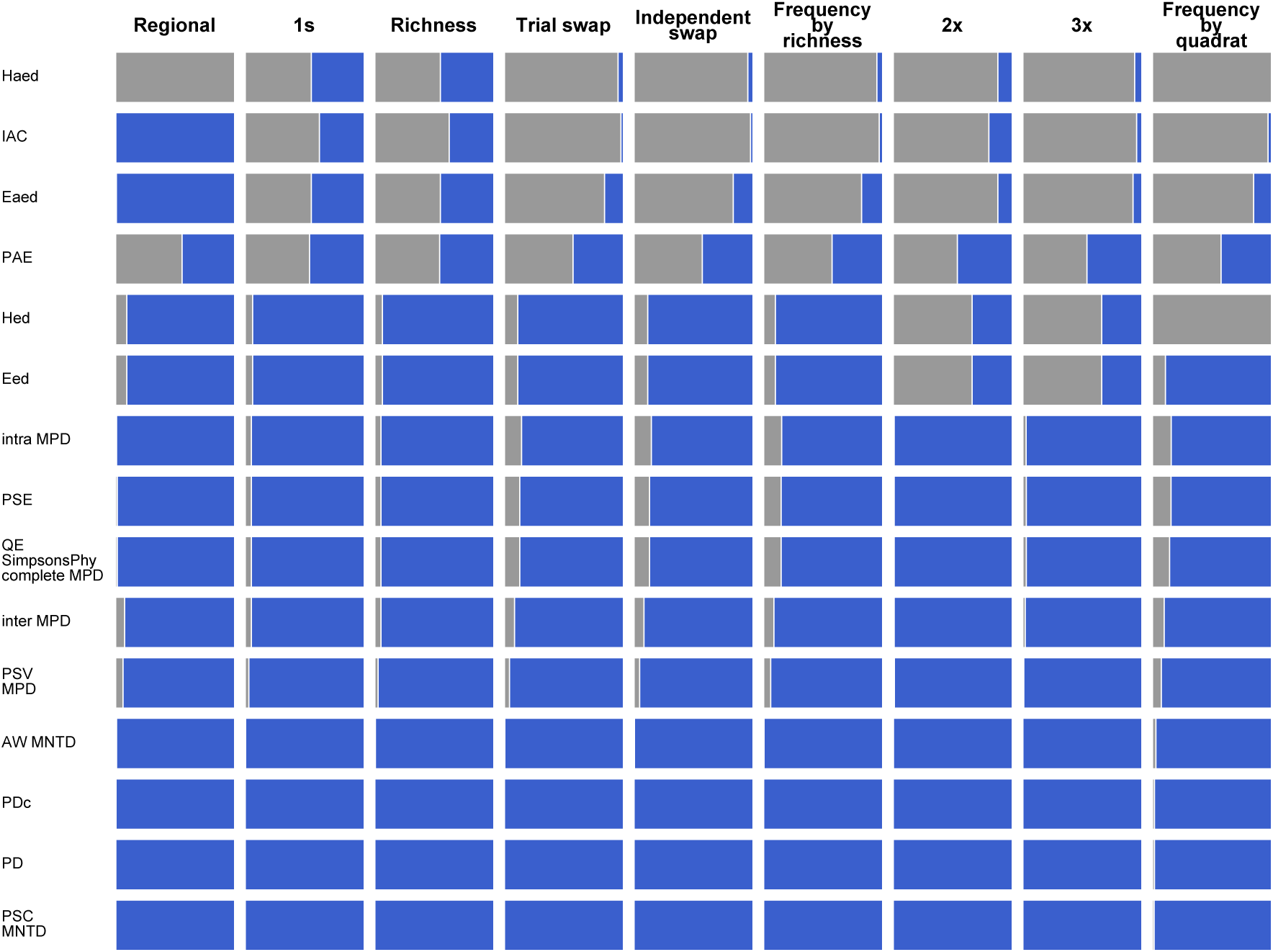
Performance of metric + null model approaches at detecting phylogenetic clustering given habitat filtering, arranged in order from best-performing to worst, with the best approaches in the bottom left corner. Blue bars summarize the proportion of the total 1,009 simulations where the mean of the standardized effect sizes was significantly less than zero (one-way Wilcoxon signed-rank test). Gray bars summarize the proportion where the mean did not differ from zero (type II errors). Equivalent metrics (e.g., PSC, MNTD) performed identically and are combined.

**Figure 4.**
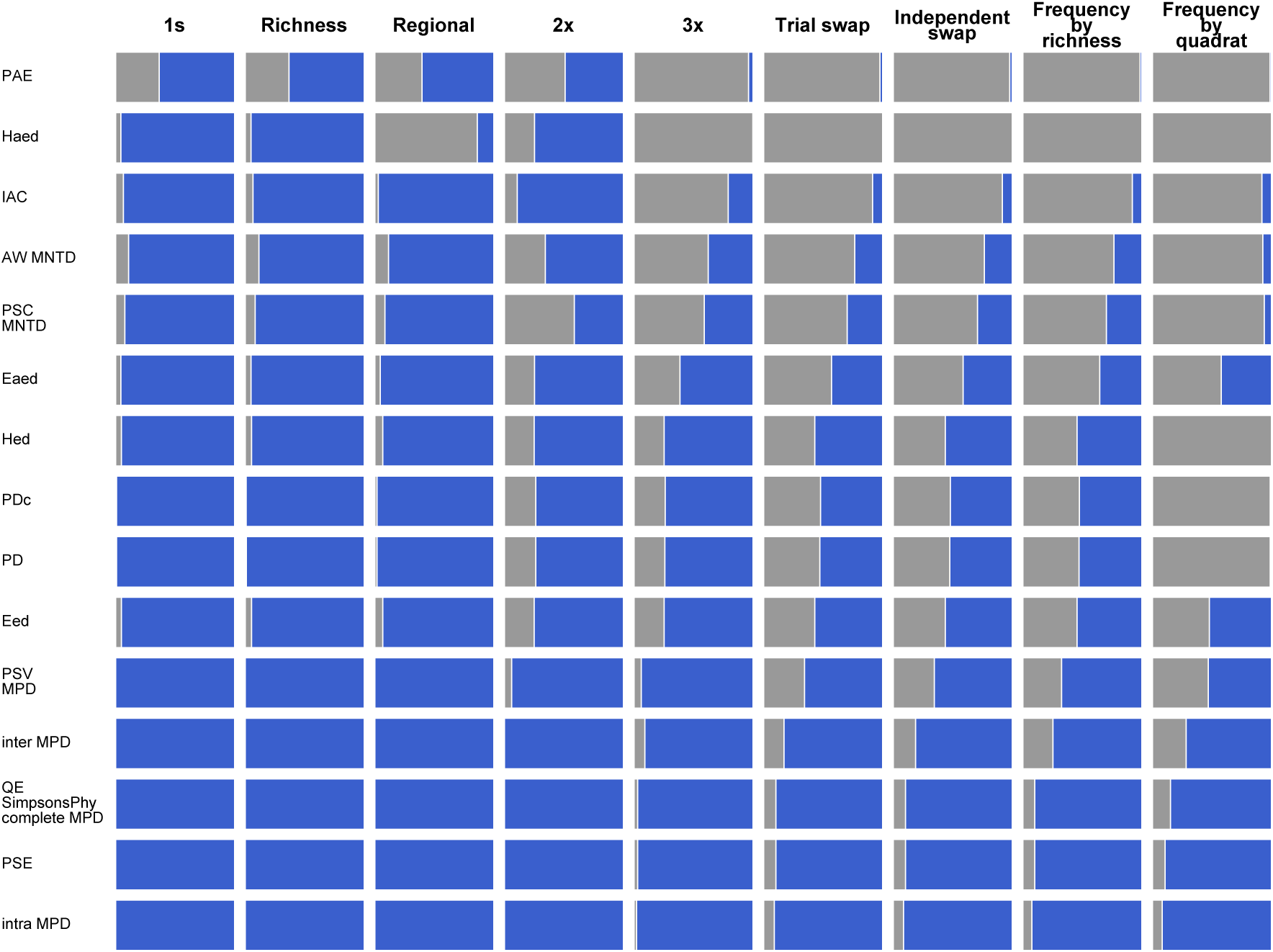
Performance of metric + null model approaches at detecting phylogenetic overdispersion given competitive exclusion, arranged in order from best-performing to worst, with the best approaches in the bottom left corner. Blue bars summarize the proportion of the total 1,009 simulations where the mean of the standardized effect sizes was significantly greater than zero (one-way Wilcoxon signed-rank test). Gray bars summarize the proportion where the mean did not differ from zero (type II errors). Equivalent metrics (e.g., PSC, MNTD) performed identically and are combined.

**Figure 5.**
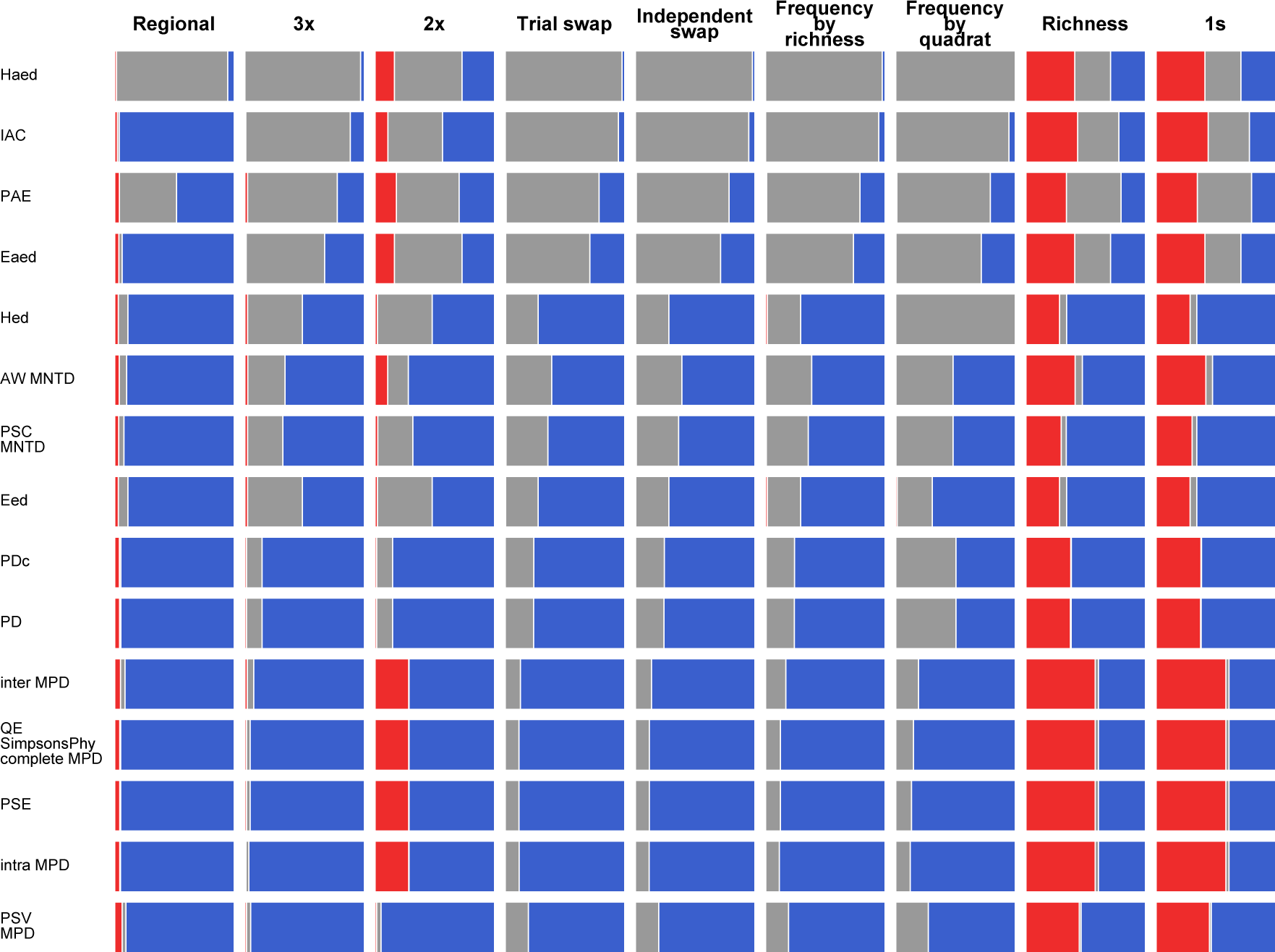
Overall performance of metric + null model approaches, arranged in order from best-performing to worst, with the best approaches in the bottom left corner. Red bars (type I errors) summarize the proportion of the total 1,009 random community assembly simulations where the mean of the standardized effect sizes differed significantly from zero (two-way Wilcoxon signed-rank test). Gray bars summarize the mean type II error rates from Figs. 3 and 4. Blue bars provide an indication of the success of each approach, and are defined as one minus the mean type I and II error rates.

## Discussion

The unification of phylogenetic community structure methods with age-old questions of community assembly has revolutionized the fields of ecology and evolution. Since Webb’s seminal papers (Webb 2000, Webb et al. 2002), there has been an explosion of interest in these matters, including a wide variety of “improvements” upon existing measures (Box 1). Many of these, however, have never been adequately tested, and others are equivalent, as we show here (Fig. 1B). Our objective was to assess a wide range of available methods in order to identify those with demonstrable utility, and to identify those that measure unique aspects of phylogenetic community structure.

Which metrics are best? The results of our study suggest that the answer depends in part on which community assembly processes are of interest, and which null models are used. However, some clear and general answers did emerge. Across most null models and all community assembly simulations, PD (Faith 1992) consistently performed well (Fig. 5), showing low type I error rates and more power than most other metrics; it was particularly good at detecting the effects of habitat filtering (Fig. 3). Group 1 (“mean relatedness”) metrics (Fig. 1) also performed well, particularly at detecting effects of competitive exclusion (Fig. 4). Like Kembel (2009), and unlike Kraft et al. (2007), we found that Group 3 (“nearest-relative”) metrics were not as powerful as Group 1 metrics at detecting competitive exclusion, though we did not directly probe changes in community size as did Kraft et al. Instead, we found that Group 3 metrics slightly outperformed Group 1 metrics at detecting habitat filtering.

We expected that because non-abundance-weighted metrics can be strongly influenced by the presence or absence of a single individual, such metrics would more frequently exhibit type I errors (Miller et al. 2013). However, abundance-weighted forms of both Group 1 metrics and MNTD showed slightly higher type I error rates than non-abundance-weighted forms. This may be because abundance-weighted metrics appear to require more randomizations before expectations stabilize (results not shown). However, increased randomizations (from 10^3^ to 10^4^) of CDMs did not alter our main conclusions (Fig. S4.1). We encourage additional exploration of the circumstances under which abundance-weighted versions of these metrics yield type I errors, and emphasize that these differences in error rates among the Group 1 metrics were small.

Some of the metrics introduced by Cadotte et al. (2010) showed poor performance, particularly PAE and *H*_AED_. The metric *E*_ED_, which out-performed other Group 3 metrics, was a notable exception, as was PD_*c*_ (though see Box 1). As suggested (Cadotte et al. 2010), these metrics do indeed measure unique aspects of phylogenetic community structure (Fig. 1B). Some of these aspects, however, do not seem to be related to traditionally recognized community assembly processes. What these metrics (PAE, *H*_AED_) quantify may yet prove useful in certain contexts (Houle et al. 2011), but they showed poor performance with the simulations in this study. When used with the regional null, IAC showed strong power to detect non-random patterns, but this did not extend to other null models. *H*_ED_ was closely correlated with PD (r = 0.94), but it did not perform as well as it. We recommend use of either PD or Group 1 metrics.

Which null models are best? Again, our results suggest that the answer depends in part on the choice of metric and the community assembly process of interest. In general, we recommend against use of a frequency by quadrat null. The CI for this null model account for neither the increased variance in expectations at smaller samples of the regional species pool (Clarke and Warwick 1998), nor the correlation of many metrics with species richness (Fig. 1). Under certain parameters (e.g., low observed quadrat species richness as compared with that of randomized quadrats), this is expected to result in high rates of type I errors, particularly for metrics that are correlated with species richness (Fig. 1A), and we suggest this null should be used with prudence.

The 2x and 3x null models showed mixed performance. While they exhibited fairly low type I error rates (Hardy 2008), they also exhibited limited power to detect expected phylogenetic community structure. When these nulls are concatenated by richness, they exhibit elevated type II error rates (Appendix S6). We suspect that extreme constraints imposed on matrix randomizations by these nulls results in biased exploration of reasonable phylogenetic space (Appendix S6). Regardless of the reason for this lack of power, the instability across species richness shown by the CI for the 2x and 3x null models (Fig. 2) means that the expectations for a given metric can change dramatically based on whether N or N+1 species are present in an observed community. Nevertheless, these null models are intended to be concatenated by quadrat, and when used in this manner, they performed better than all but the regional null.

The regional null (Appendix S1) was designed to simulate dispersal, proportional to species abundance in a regional pool, into a local community (study area) of interest, such that deviations from these dispersal pressures (e.g., the product of environmental filters) can be readily detected, and local community dynamics (e.g., competition) do not obfuscate expectations. For instance, given strong competitive exclusion, local communities may show widespread phylogenetic overdispersion, where certain species are generally excluded. When these observed occurrence frequencies are taken as regional occurrence frequencies and randomized accordingly (as in the independent swap), it becomes difficult to detect phylogenetic overdispersion, since the randomized CDMs will tend to contain distantly related species, and confidence intervals are accordingly shifted up from those expected given a model like the richness null (Fig. S1.6). The regional null avoids this issue by using expectations from a larger, fixed pool as the standard against which to compare observations from the study area. However, it is difficult to quantify dispersal pressure on a community of interest, and this model may not be practical for many researchers. Future studies should investigate what information might be used to construct these expectations (e.g., range sizes), and whether this null can be of widespread utility (Lessard et al. 2012).

Null model choice cannot be driven entirely by statistical properties. There may be sound biological reasons for why a given null should be employed, even if its statistical performance is not on par with others (Gotelli and Graves 1996). However, such reasoning should not come at the expense of common sense. For instance, if the quadrats from a CDM are not thought to be representative of the study area (e.g., biased sampling across study areas), then a null model like the independent swap that maintains these observed occurrence frequencies will only confuse interpretation of results. In short, we recommend use of a model that randomizes data structures relevant to the hypothesis, while maintaining structures unrelated to that hypothesis, and while being cognizant of null model performance (Figs. 3-5) and behavior (Fig. 2). The behavior of any metric + null approach with any CDM can be elucidated with use of the expectations function from *metricTester*. More recently, efficient algorithms for directly calculating the richness-standardized forms of MPD and PD (i.e. SES after randomization with richness null) have been developed that do not require lengthy randomizations (Tsirogiannis and Sandel 2015), and there is room to extend such an approach to additional metrics and nulls.

What combined approach do we suggest? The richness null with PD or Group 1 metrics may offer the simplest results to interpret by making the clearest assumptions (any species can occur anywhere); more constrained null models raise questions of sampling artifacts and the efficiency of swap algorithms. We emphasize that little should be made of the deviation of any single CDM beyond expectations; the high type I error rates of most approaches casts doubt on the interpretation of single community tests. However, if a metric that is uncorrelated with species richness is used (e.g., PSV), then quadrats from that CDM can be arranged along an environmental gradient to test hypotheses (Graham et al. 2009, Miller et al. 2013). Here, the slope is of interest, rather than the significance of any individual community (e.g., quadrats are increasingly phylogenetically clustered along a gradient of decreasing precipitation). Hypothesis testing in this manner minimizes the necessity of a null model, and raw metric values, which often have intrinsic meaning, can then be used instead of SES. For instance, the MPD of a community, given a time-calibrated phylogeny, is equal to the mean evolutionary time separating co-occurring taxa. Other metrics like PD are correlated with species richness, and should be used with a null model (or otherwise standardized, e.g., Nipperess and Matsen 2013) if the focus is on phylogenetic community structure (as opposed to e.g., PD itself). Researchers need to consider what they are measuring with their metric(s) of choice, whether they need to standardize those metrics, and why or why not they might procure significant results.

By making the assumption that the traits responsible for community assembly covary with phylogeny, this study maintains the sometimes questionable dogma that habitat filtering leads to phylogenetic clustering, and that competitive exclusion leads to phylogenetic overdispersion (Webb et al. 2002, Mayfield and Levine 2010). If trait data are available, we encourage researchers who use these methods to fit explicit models of evolution to traits pertinent to the assembly processes in question (Butler and King 2004), and to also investigate patterns of community structure in functional traits. In this study we did not test approaches that account for variation among quadrats in species co-occurrence probabilities (e.g., Cavender-Bares et al. 2004; Hardy & Senterre 2007), but *metricTester* could be adapted to investigate these metrics. There is also an expansive assortment of existing (and yet to be created), hypothetically useful null models whose behavior and performance remains to be tested (e.g., Ulrich and Gotelli 2010). Ultimately, advanced approaches (Ives and Helmus 2011) may prove more powerful and gain wider use than current phylogenetic community structure metrics, but the existing arsenal remains well suited to addressing a wide variety of questions.

## Acknowledgements

We thank Vincenzo Ellis, Matt Pennell, Amy Zanne and the Ricklefs lab for input and feedback, the Harmon lab for help with coding, and the Oslo Bioportal, the University of Missouri Lewis Cluster, and the Domino Data Lab for providing computing resources. We thank Alex Kharbush for providing logistical and technical advice with Amazon Web Services.

## Data accessibility

*metricTester* is available from GitHub (<u>https://github.com/eliotmiller/metricTester</u>), and can be directly installed into an active R session using the *devtools* package. It requires the package *ecoPDcorr*, which can also be directly installed with *devtools* (<u>https://github.com/eliotmiller/ecoPDcorr</u>).

## Appendix S1

Null models: behavior across variation in species richness, documenting equivalency, and regional null model description.

### Behavior of existing null models across species richness

As described in the main text, we were interested in quantifying the behavior of the null models (Table 2) across varying species richness. Basic principles of bootstrapping (Efron 1979) suggest that there should be more variance when small subsamples of a larger pool are taken. If two random taxa are drawn from a phylogeny, they could be close sister species, or they could span the root. The calculated phylogenetic community structure metrics from these two extremes would vary greatly. Alternatively, if all the species from a phylogeny are present in a community, we know what the calculated metric will be (+/- some slight variation for abundance-weighted metrics). This should lead to a confidence funnel (Clarke and Warwick 1998), with more variable expectations at lower species richness. But what sorts of expectations do the different nulls we tested generate? How do they differ from each other? What factors influence their distributions?

The richness null (=SIM3, Gotelli (2000)) we tested swaps abundances within quadrats. In other words, given a quadrat by species community data matrix (CDM), this null shuffles the contents of each row (a quadrat). Accordingly, each species is sampled with equal frequency in the randomized matrices (i.e. it does not maintain species’ observed occurrence frequencies). We would expect that for metrics like mean pairwise phylogenetic distance (MPD) that are uncorrelated with species richness (Fig. 1), the mean expected value would not change with species richness. Simulations show this is the case (Fig. 2). This is a useful null to use as a benchmark against which to understand other more constrained nulls. A slight variation on this, the 1s null model (Hardy 2008), converges on the same expectations as the richness null (Fig. S1.1). The 2s null (Hardy 2008) is Hardy’s implementation of the richness null, and we examined it here simply to confirm that different R packages do indeed give similar solutions (Fig. S1.1).

The frequency null we tested (=SIM2, Gotelli (2000)) swaps abundances within species. We refer to this as the frequency by quadrat null. Given a quadrat by species CDM, this null shuffles the contents of each column (a species). This means that individual species are not sampled with equal frequency. Importantly, it also means that the randomized quadrats tend to contain the mean number of species as were observed in the input CDM. For example, given a CDM with four quadrats, one of species richness 2, two of species richness 5, and one of species richness 8, randomized quadrats will tend to contain 5 species. Based on the principles of bootstrapping mentioned above, it should be clear why this would be problematic; the larger expected variance at low species richness will not be incorporated in the null model, and high type I error rates are expected (black arrow in Fig. 1 points to the region of concern).

To account for this, Miller et al. (2013) developed a method where the per quadrat raw metric values and associated species richness from a frequency null were retained. These values were concatenated by species richness, and observed values were compared to those expected at their corresponding species richness. We refer to this as the frequency by richness null.

Like the frequency by richness null, the derivation of a CDM where the randomized quadrats contain the same number of species as the input CDM, and individual species occur with the same frequency as the input CDM are the goals of the independent swap (Gotelli and Entsminger 2001) and trial swap null models (Miklós and Podani 2004). The trial swap null model has been considered a more efficient implementation of the independent swap (Miklós and Podani 2004). In our simulations this was not the case (although we are not entirely sure whether convergence on stable confidence intervals is necessarily equivalent to the “equidistribution” of randomized matrices discussed by those authors). With increasing randomizations of a given CDM, the independent swap, trial swap and frequency by richness nulls all show increasingly stable expectations, but the trial swap seems to stabilize at a slower rate (Fig. S1.2). Regardless of the reason for this result, all three nulls converge on the same solution (Figs. S1.3 and S1.4).

The 2x and 3x nulls (Hardy 2008) were developed to maintain not only aspects of species richness and occurrence frequency, but also either the quadrat-specific rank abundance curve or the species-specific abundance distribution, respectively. While these are aspects of a dataset that a researcher most certainly might wish to maintain, in practice, the extreme constraints imposed on the matrix randomizations seems to result in inefficient exploration of phylogenetic space (Appendix S6). Both nulls also gave identical solutions when concatenated by richness (Fig. S1.5). We were unable to determine why these nulls behaved as they did, but the fact that their expectations wobble across species richness seems to be an undesirable property. Biologically, it is hard to construct a reason why one should expect dramatically different phylogenetic community structures with the presence or absence of a single species. Indeed, when using a non-abundance-weighted metric like MPD, a null model that maintains species richness and occurrence frequency such as the independent swap and a null model that maintains both of these aspects of the data and aspects of abundance distributions like the 2x and 3x null models should converge on similar expectations. Using MPD, quadrat-specific and species-specific abundance distributions should not influence the expectations from two quadrats of similar species richness (e.g., there is no reason to expect the randomized values from two observed quadrats, each with five species, to converge on different expectations). Instead, quadrats of the same species richness converged on different expectations, suggesting poor exploration of possible phylogenetic community space (Appendix S6). Concatenating expectations from quadrats of the same species richness resulted in odd-shaped randomized distributions and a total loss of power to detect simulated community assembly processes (Appendix S6). That said, when concatenated by quadrat, as they were intended to be, these null models did perform better than most we tested (Fig. 5).

What determines how expectations for the independent swap (or frequency by richness or trial swap) vary from those given the richness null? It may not be intuitive to all readers that species within a phylogeny vary in their mean phylogenetic distance to other species in the phylogeny. In an ultrametric tree, all species are equidistant from the root. How can one differ from another in its mean relatedness to other species? Consider the case of a single species that is sister to the rest of the phylogeny. This species is separated by larger average evolutionary distances than are the other species. The relationship between species’ occurrence frequencies and their mean relatedness determines how the expectations for the independent swap shift from those of the richness null.

To illustrate this point, we generated a CDM as described in the main text. For every species in the CDM, we next calculated both its mean relatedness to the rest of the species and its occurrence frequency in the CDM. In the first simulation (Fig. S1.6, “sim1”), we then replaced species identities in the CDM such that species that were more closely related to the rest became the most frequently occurring species in the CDM. In other words, the most closely related species in the phylogeny also became the most common in the new CDM. We performed the opposite procedure in the second simulation (“sim2”). When distant relatives are also the least frequently observed species, the expectations are shifted downwards from those given a richness null. When distant relatives are the most frequently observed species, the expectations are shifted upwards (Fig. S1.6). Moreover, mean expected MPD, which is uncorrelated with species richness, begins to show some correlation with species richness when using a null model like this. This is because the probability of including rare species in the randomized matrices increases with larger samples. Thus, the expected MPD is positively correlated with species richness in the first simulation, and negatively correlated in the second.

### Development of the regional null model

No null model of community assembly that we know of maintains species richness, species occurrence frequency, and species abundance. The null models that come closest to achieving these objectives are the 2x and 3x nulls of (Hardy 2008). We developed a regional null model aimed at achieving these goals. We did this in particular because our competitive exclusion simulations led us to recognize the importance of local interactions on species occurrence frequencies (Appendix S3), though the importance of considering the regional pool has also been recognized widely in the literature (Ricklefs 1987, Lessard et al. 2011, 2012). Specifically, our competitive exclusion simulations produce a local effect where some species that are abundant in the regional species pool become locally less so (Fig. S4.6). Such species are closely related to species that are more abundant in the local community (i.e. the simulation arena and resulting CDM). When these local occurrence frequencies are used to inform a null model such as the independent swap, short phylogenetic distances (like those between sister species) tend not to occur in the randomized matrices, which results in the expected phylogenetic community structure being shifted upwards from that given a null that maintains only species richness (Fig. S1.7). Accordingly, it becomes difficult to detect phylogenetic overdispersion.

In empirical situations, researchers are likely interested in testing for the effects of community assembly processes in a focused area (e.g., a forest plot, a grid cell on a map, a soil sample, etc.). The thought, likely, is that the focal area was historically or is currently subject to community assembly processes (e.g., competitive exclusion) that operate on a smaller scale than regional dispersal dynamics. The regional null is intended to simulate these regional dispersal probabilities into the focal area. The regional null model largely accomplishes the objectives of maintaining species richness, occurrence frequency, and abundance distributions, and was associated with lower error rates than the other null models (Fig. 5). It requires, however, that a regional abundance vector (in the form of “sp1, sp1, sp1, sp2, sp2, …”) be provided. Developing a vector like this is easy in our simulations, but may be more difficult in empirical situations. If a dataset consisted of evenly sampled sites, so as not to introduce biases in species occurrence frequencies, and the assumption was made that species abundances reflected their dispersal probability, then a vector of all individuals across the entire dataset could be used (use the function abundanceVector in *metricTester* to do so). Most real-world situations are more complicated than this, and the practicality of the regional null remains to be demonstrated.

The regional null takes as input a regional abundance vector and, for each quadrat in the randomized CDM, it then samples with equal probability from this vector the same number of individuals as were in the given quadrat in the observed CDM. The metric of interest is calculated on the quadrats from this randomized CDM, and these values are retained, along with the associated species richness from each quadrat. This process is repeated many times. The randomized values are then concatenated by their associated species richness. Thus, species richness is strictly maintained, as observed quadrats are only ever compared with randomized sites of corresponding species richness.

Species occurrence frequencies are also approximately maintained with the regional null. For instance, after 1,000 randomized CDMs were generated with the regional null, we calculated the mean occurrence frequency across all randomized CDMs for each of the 50 species in CDM. These values were closely correlated with the observed occurrence frequencies for the same 50 species (*r*^2^ = 0.83, *p* < 0.001, Fig. S1.8). The abundance at which a species occurs in any given quadrat is also approximately maintained with the regional null. For instance, within a given quadrat from these same randomizations, a randomly selected species was mostly found as a single individual, occasionally as two individuals, and very infrequently at higher abundances (Fig. S1.9A). This is similar to the abundance distribution of the same species in the original CDM (Fig. S1.9B).

**Figure S1.1.**
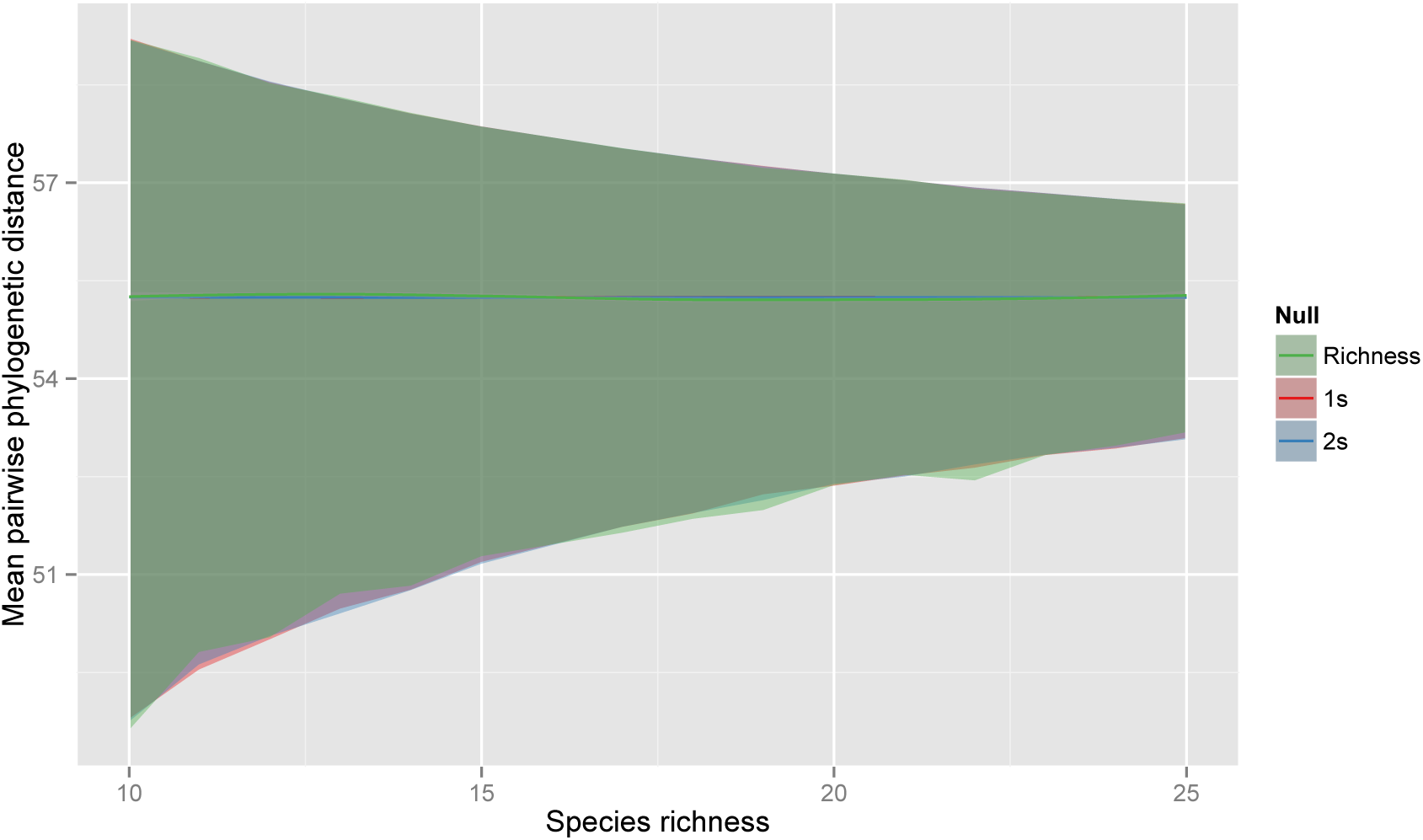
Confidence intervals (95%) for null models (shaded by color) across variation in species richness. The same initial CDM, phylogeny and number of randomizations as Fig. 1 were used. The richness and 1s null models provide identical expectations. The 2s null model also converges on the same expectation; this model is simply the *spacodiR* implementation of what amounts to a richness null, but we include it here to confirm that different R packages provide similar results.

**Figure S1.2.**
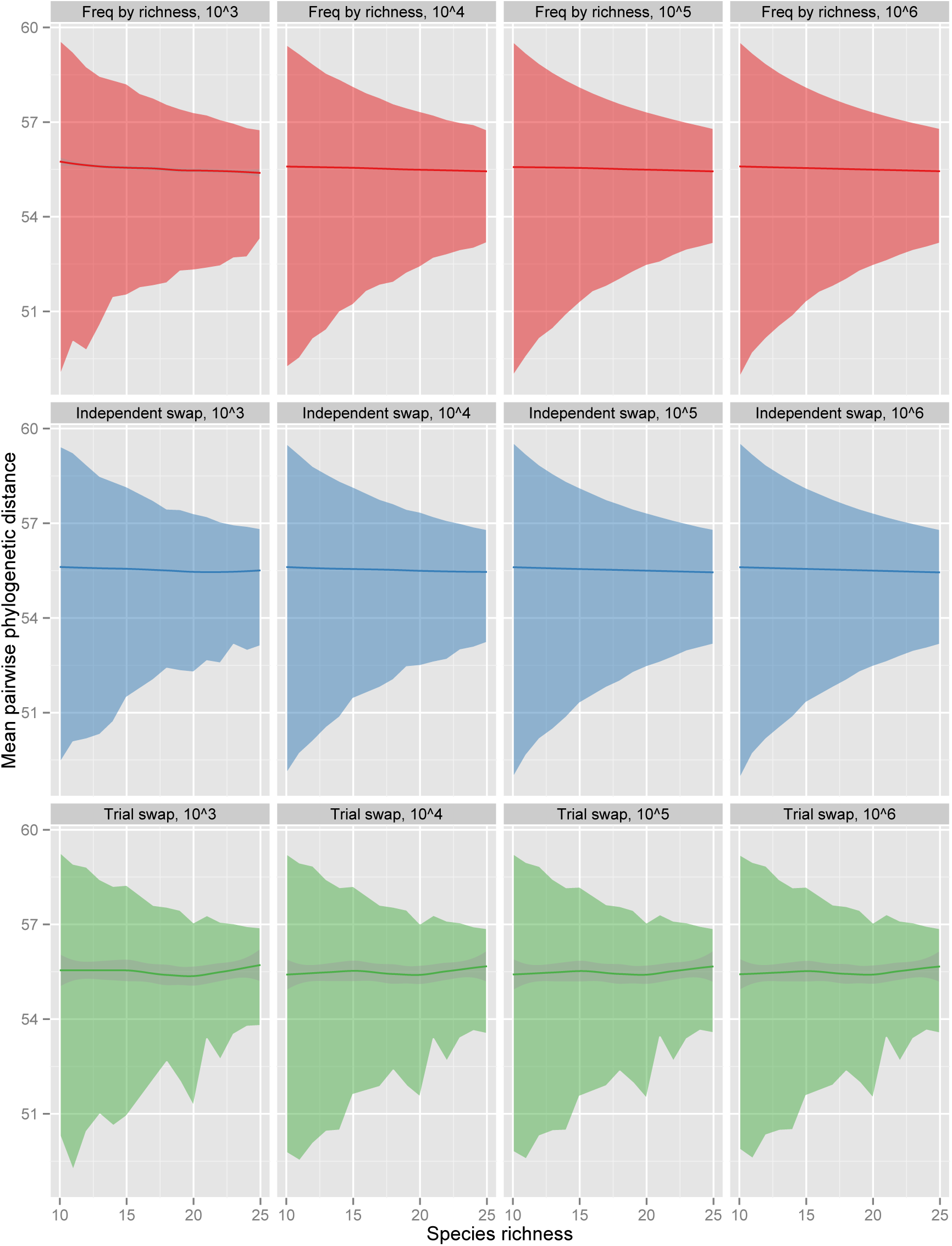
Confidence intervals (95%) for the frequency by richness, independent swap, and trial swap nulls across varying species richness and with increasing randomizations of an initial CDM. The darker lines in all panels represent mean trend lines. The shading around those lines represents confidence around the mean, though shading is not visible (in the first two models) where there is a great deal of confidence around the mean.

**Figure S1.3.**
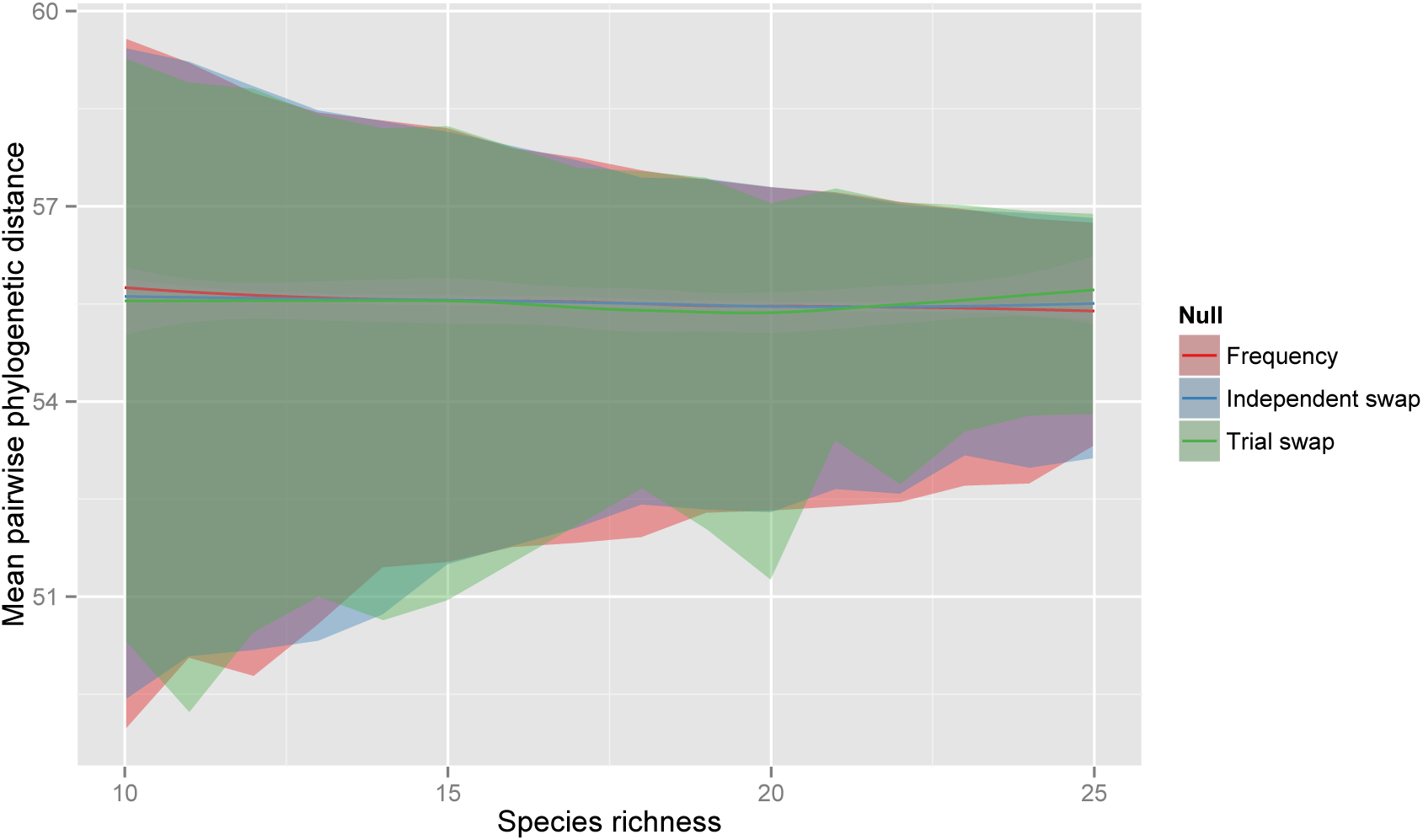
Confidence intervals (95%) for the frequency by richness, independent swap, and trial swap nulls across species richness (after 10^3^ randomizations). These are the leftmost three panels from Fig. S1.2. Darker lines represent mean trends. Shading around those lines represents confidence around the mean; the shading is only visible for the trial swap mean.

**Figure S1.4.**
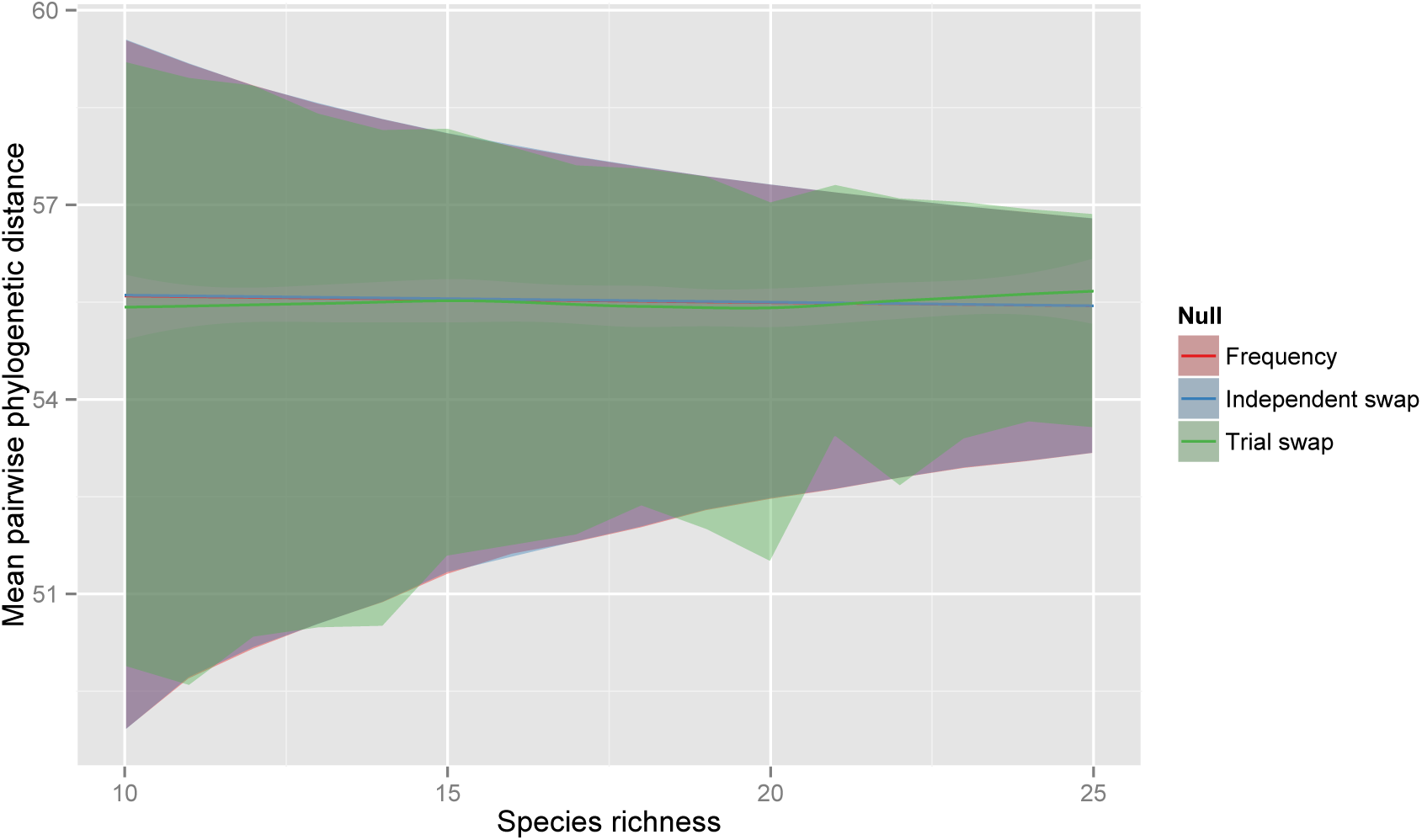
Confidence intervals (95%) for the frequency by richness, independent swap, and trial swap nulls across species richness (after 10^6^ randomizations). These are the rightmost three panels from Fig. S1.2. Darker lines represent mean trends. Shading around those lines represents confidence around the mean; the shading is only visible for the trial swap mean.

**Figure S1.5.**
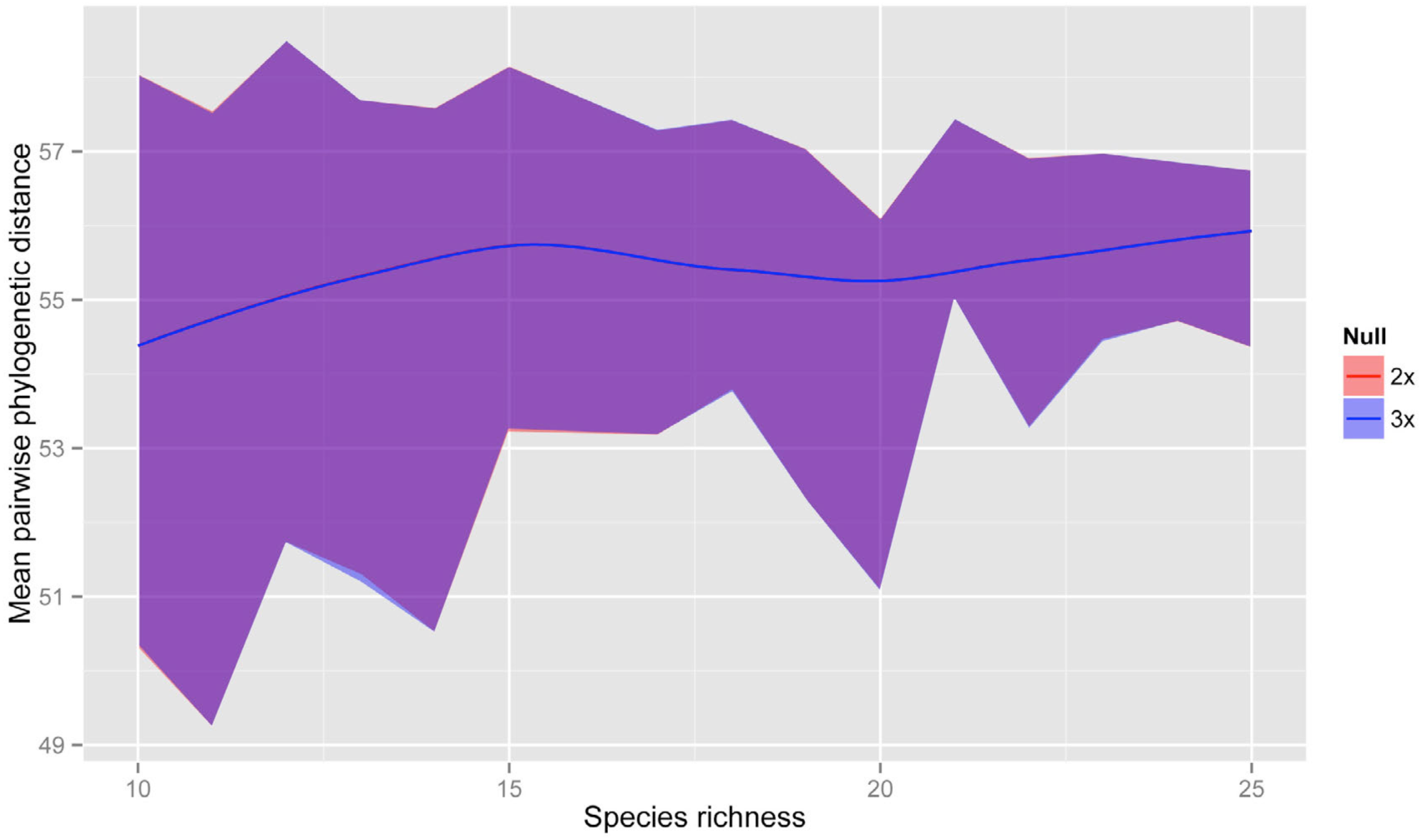
Confidence intervals (95%) for the 2x and 3x null models (Table 2) across variation in species richness. Expectations shown here are the result of 10^5^ randomizations. The two null models follow identical distributions.

**Figure S1.6.**
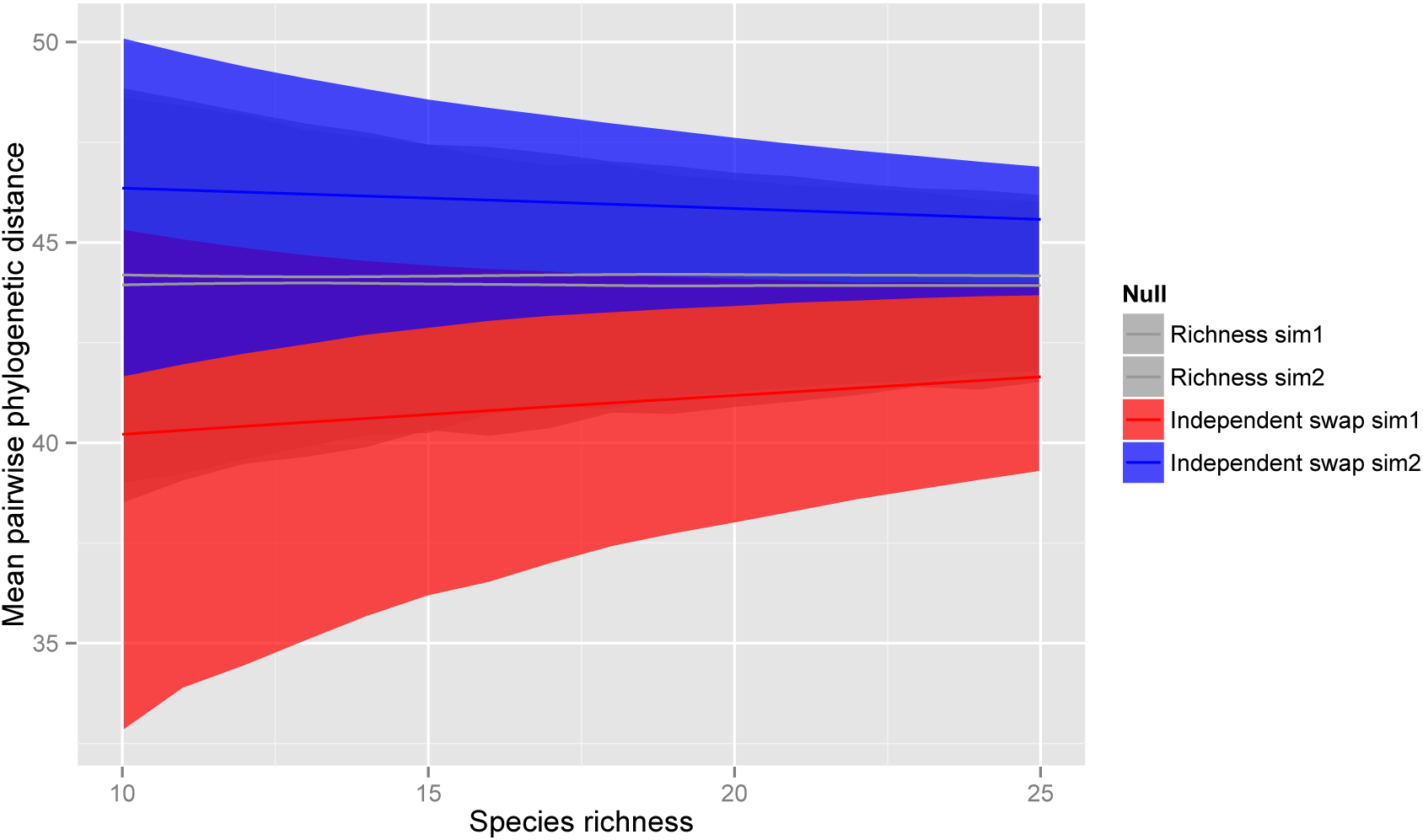
Results of two simulations varying the occurrence frequency of individual species in the CDM according to mean relatedness to the rest of the species in the phylogeny. In the first simulation, the most closely related species occurred most frequently. This pattern was reversed in the second simulation. Expectations do not shift notably when using the richness null.

**Figure S1.7.**
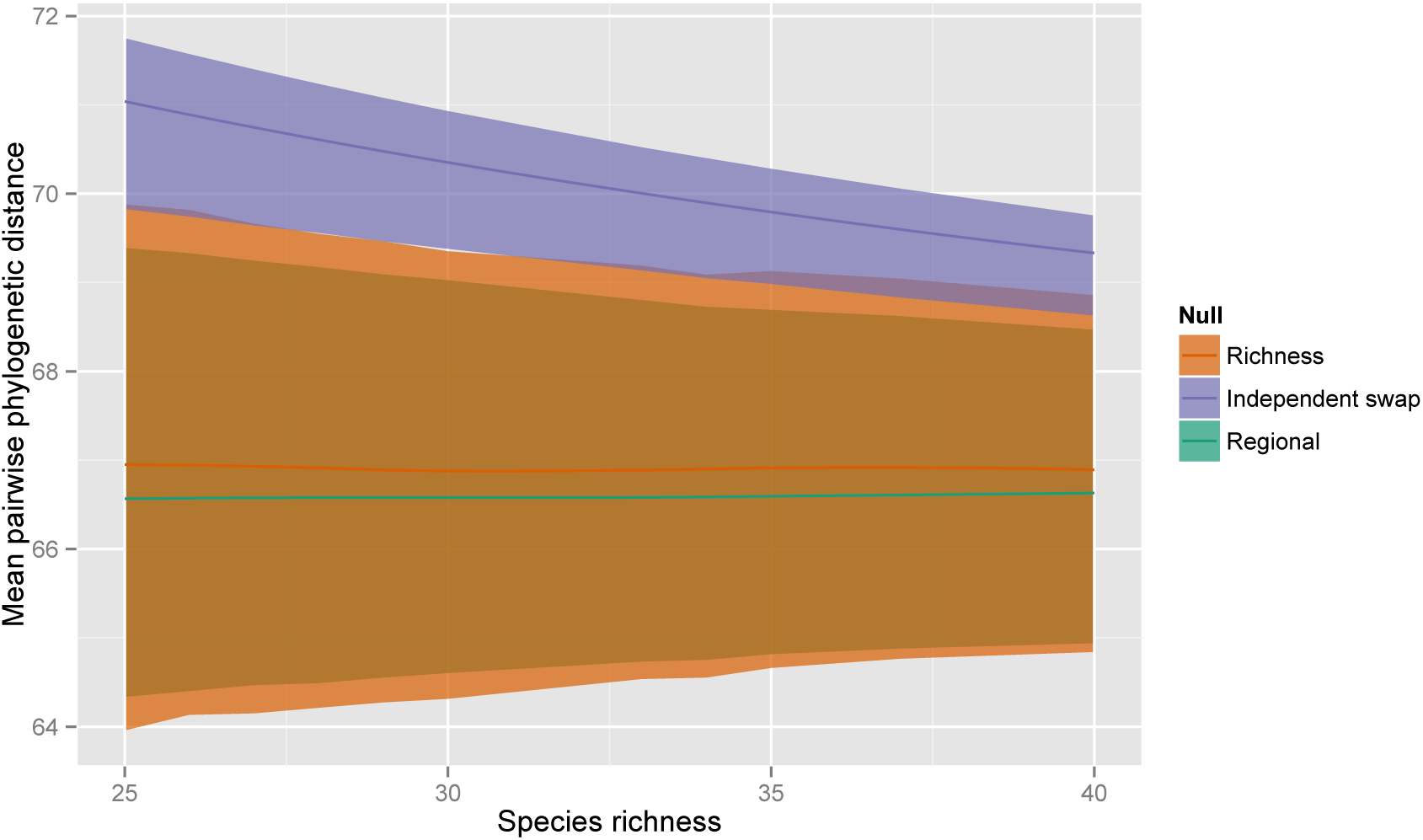
Confidence intervals (95%) for null models (shaded by color) across varying species richness. The arenas were constructed with the same parameters as described for the competitive exclusion simulations in the main text. The expectations shown here are the result of 10^5^ randomizations. After multiple generations of competition, the abundance of some closely related species (i.e. species more nested in the phylogeny) decreases across the arena (Fig S3.11), as does their occurrence frequency in the random quadrats, used to generate the CDM. Thus, the expectations given an independent swap null, which accounts for occurrence frequency, are shifted notably upwards from those given a richness null. Moreover, some species are lost from the arena entirely, and the mean expectations for the richness null are therefore also shifted slightly up from those given the regional null.

**Figure S1.8.**
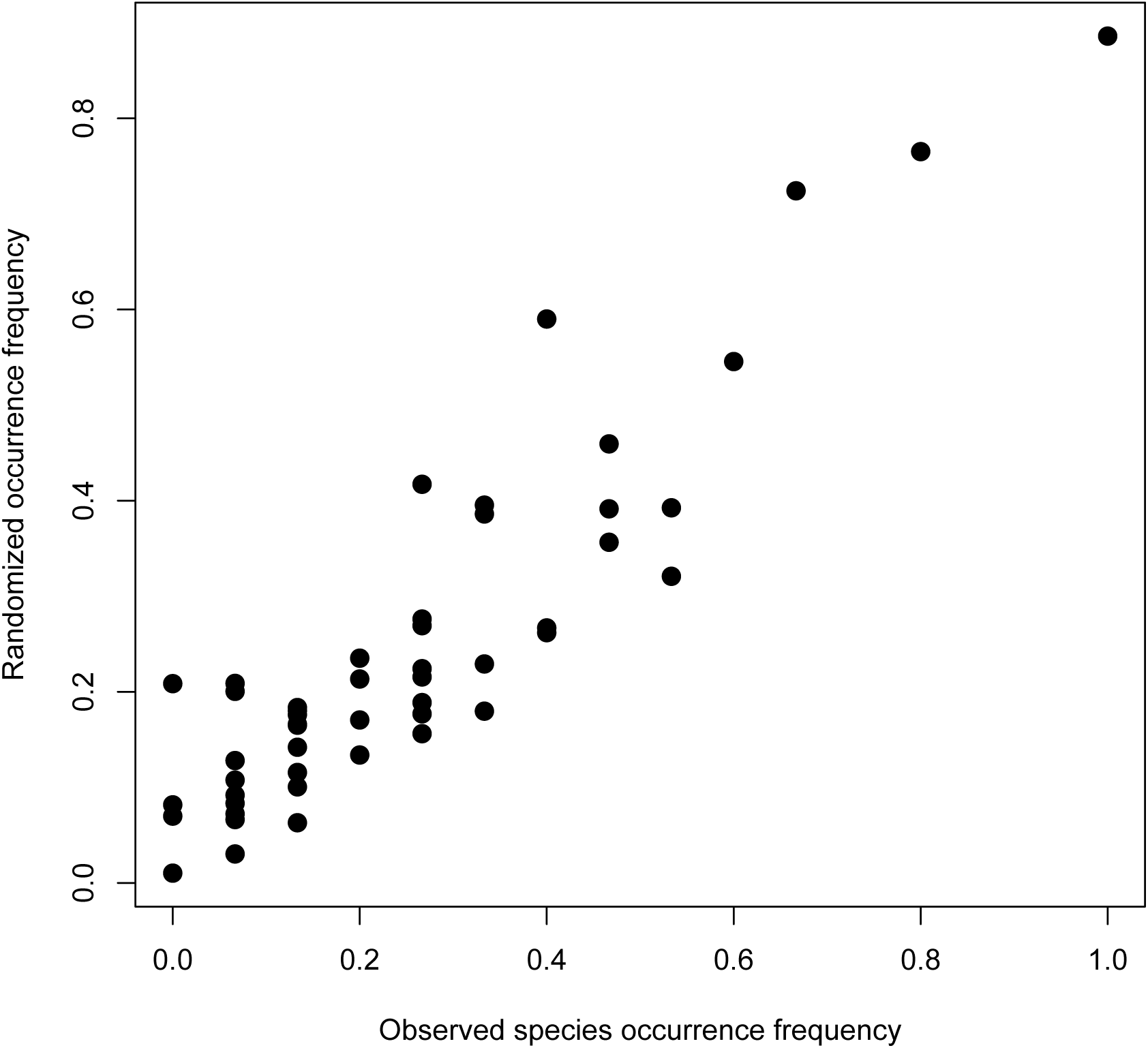
Mean occurrence frequency of 50 species, after 1000 randomizations with the regional null model, as compared with their initial occurrence frequency. Species tended to occur with a frequency proportional to their occurrence frequency in the observed matrix (*r*^2^ = 0.83, *p* < 0.001).

**Figure S1.9.**
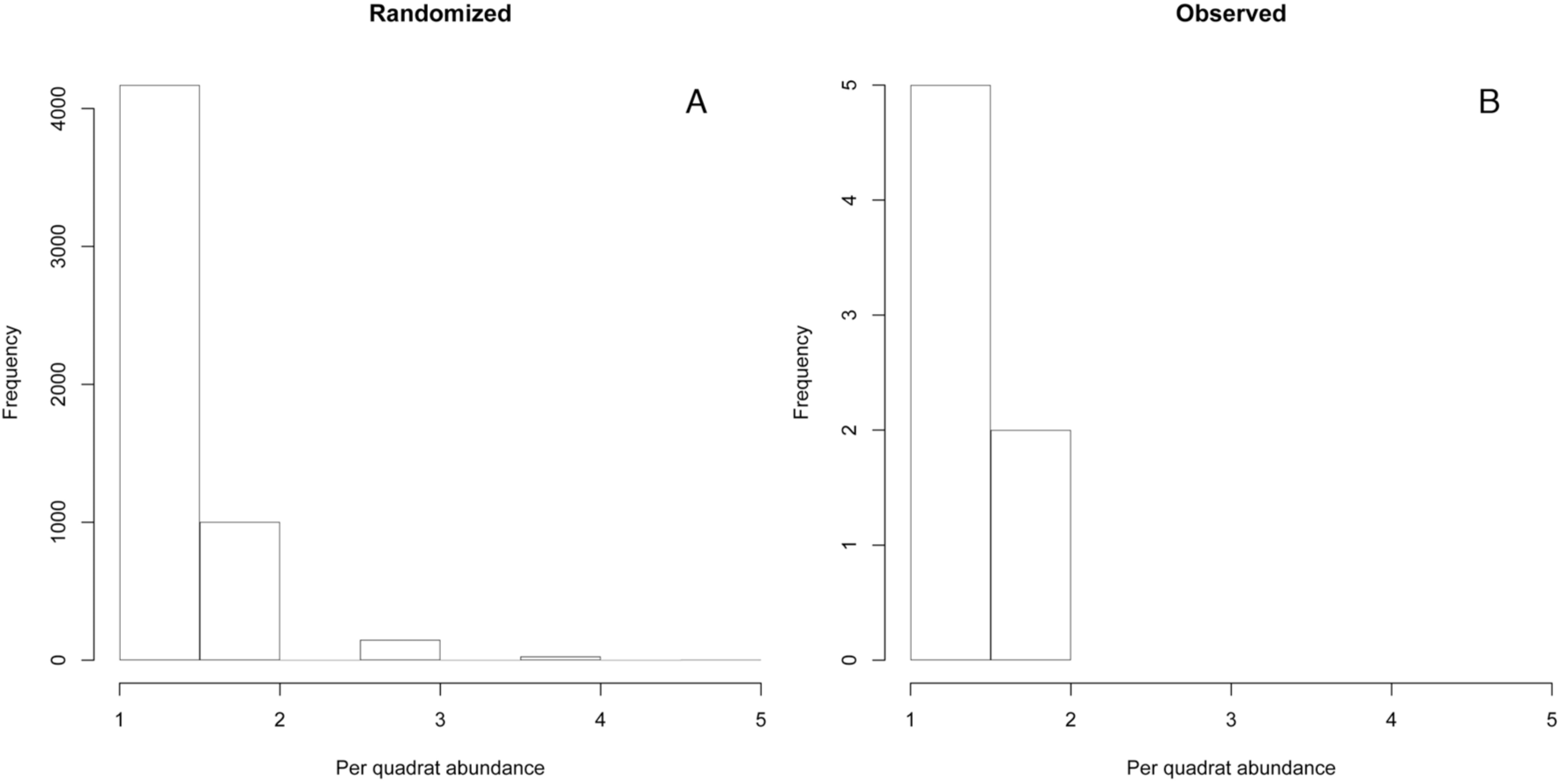
(**A**) Histogram of the abundance distribution of a randomly selected species across 1000 randomized community data matrices (after excluding all quadrats where it did not occur at all). (**B**) Histogram of the observed, original abundance distribution of the same randomly selected species as **A** (after excluding all quadrats where it did not occur at all).

## Appendix S2

Three forms of abundance-weighted MPD, and equivalency of some forms to Clarke and Warwick’s metrics.

### Three forms of abundance-weighted MPD

Abundance-weighted mean pairwise phylogenetic distance (MPD) and mean nearest taxon distance (MNTD) were introduced in *Phylocom* (Webb et al. 2008) without accompanying publications. These methods have entered into common usage in the literature, but they have not been discussed at any length. A variation on abundance-weighted MPD was recently introduced that only accounts for interspecific phylogenetic distances (Miller et al. 2013). This is different than the implementation in *Phylocom* and *picante* (Kembel 2009).

There are at least three different possible forms of abundance-weighted MPD (Fig. S2.1). Consider a local assemblage of three species drawn from a regional species pool. Qualitatively, species A, B, and C are clustered in the phylogeny. But, how should the abundances of these three species affect the metric? In the simple case of an assemblage of two individuals of species A, and one each of species B and C, all of the potential interactions among individuals can be visualized schematically (Fig. S2.1).

If we include only interactions among heterospecific individuals to derive a matrix of abundance weights for the MPD calculation (Fig. S2.1, “interspecific”), we obtain the MPD among heterospecific individuals within the community. This is the same as the MPD among species, weighted by the number of individuals of each interacting species. It is also the same as Δ* of Clarke & Warwick (1998) (see below). The resulting MPD calculated with this metric is slightly less than the unweighted version. This slight decrease is due to down-weighting in the calculation of the contribution of the phylogenetic distance between individuals of the rarer species, B and C, compared to that of unweighted MPD (Fig. S2.1).

The interspecific metric will be useful when it is the phylogenetic distances among individuals of different species that are of interest. For example, when testing for habitat filtering or interspecific competition, given an increase in the number of individuals of species A, a researcher might prefer not to have the metric show a dramatic increase in the degree of clustering (as happens with alternative versions of the metric, see below and Fig. S2.2d). This is because it is the phylogenetic distances among individuals of different species that are hypothesized to be clustered and/or overdispersed. As another example, a researcher studying phylogenetic niche conservatism might be interested in how phylogenetic community structure changes along an environmental gradient. Given abundance data, he or she could study these changes along the gradient, down-weighting the importance of rarely recorded species (e.g., vagrants) and up-weighting the importance of abundant species.

Alternatively, one might wish to account for both inter-and intraspecific interactions to obtain the mean pairwise phylogenetic distance between any two individuals within the community (Fig. S2.1, “intraspecific”). Here, the two intraspecific interactions for species A, which correspond to phylogenetic distances of zero, are given weight when calculating MPD, considerably decreasing the resulting metric from the unweighted version. This intraspecific abundance-weighted MPD is equal to Δ of Clarke & Warwick (1998) (see below). It will likely be preferred when examining patterns in community phylogenetic structure predicted to arise from processes generating negative density-dependence mediated by phylogenetic relatedness. For example, in the case of pathogen-mediated species co-occurrence, the inclusion of both intra-and interspecific phylogenetic distances is important as both con-and heterospecific individuals represent potential hosts, and the expectation may be not only of even spacing among species, but even abundance distributions of individuals among species.

Lastly, abundance-weighted MPD, as currently implemented in *Phylocom* and *picante*, is calculated by accounting for all possible interactions, including those of an individual with itself (Fig. S2.1, “complete”) (Webb et al. 2008, Kembel et al. 2010). The biological interpretation of this metric seems more complicated than those of the interspecific or intraspecific methods. The complete method might be likened, biologically, to including an individual’s impact both on others and on itself; for example, an individual’s use of environmental resources reducing availability for all individuals, including itself. The diagonal element in the abundance weight matrix of the complete method is equal to *n*^2^, where *n* is the number of individuals of a species, while that in the intraspecific method is *n*^2^ – *n*. Thus, MPD values calculated with either the intraspecific or complete versions will converge rapidly as *n* increases (Fig. S2.3). Only at low total local assemblage abundance is the difference in MPD values between these metrics notable. Nevertheless, it seems that intraspecific MPD is a more accurate implementation of abundance-weighted MPD as defined by Webb *et al*. (2008) to be the average phylogenetic distance between any *two individuals* drawn from a sample.

Each of these methods corresponds to a different biological interpretation, and they performed similarly overall (Fig. 3-5). A few points should still be understood about the intraspecific and complete methods. Both intraspecific and complete abundance-weighted MPD will correlate with assemblage species richness, since at lower richness, proportionally more intraspecific phylogenetic distances (i.e. distances of zero) are included in the mean (Fig. 1). Also, assemblages of uniform species abundances will have different MPD scores depending on whether they are abundance-weighted or not (Fig. S2.2). Finally, abundance-weighted MPD will always be less than the unweighted form (except in the unique case where all species in the assemblage are represented by a single individual, Fig. S2.2).

It is instructive to consider how these three different MPD metrics change as species abundances vary. If all species’ abundances are increased, keeping relative abundances the same, the resulting metric is unchanged for the interspecific and complete methods, but decreases for the intraspecific method (it converges on the complete method with increasing total assemblage abundance, Fig. S2.3). If individuals of both species A and C are increased in tandem towards infinity, holding B constant, then the interspecific method converges on the phylogenetic distance between species A and C (4 in this example), while the latter two methods converge on the mean of the phylogenetic distance between species A and C and their intraspecific phylogenetic distance (2 in this example; the mean of 4 and zero). Similarly, with the interspecific method, adding individuals of species A only to the assemblage will increase the contribution of the phylogenetic distances between species A and other species, while with either of the other two methods, it will increase the contribution of both interspecific distances involving species A, and distances within species A (Fig. S2.2).

*Some forms of MPD are equivalent to Clarke and Warwick’s earlier metrics*

While writing this manuscript, we became aware of three additional phylogenetic community structure metrics that were not incorporated in the main simulations (Clarke and Warwick 1998). This oversight was due in large part to the fact that these metrics have been more frequently used by conservation biologists than by community ecologists (Box 1). As we show here, they are equivalent to other metrics that we did assess, and consequently are expected to perform equivalently. Specifically, non-abundance-weighted MPD is equal to Δ+, interspecific MPD is equal to Δ*, and intraspecific MPD is equal to Δ (Fig. S2.4).

R code to demonstrate the equivalency of the metrics is provided below. *metricTester* can be installed directly from GitHub using the *devtools* package (username = “eliotmiller”; note that the dependency *ecoPDcorr* must also be installed using the same username).

~~~
library(metricTester)

library(geiger)

#simulate tree with birth-death process

tree <-sim.bdtree(b=0.1, d=0, stop=“taxa”, n=50)

#generate log-normal abundance curve

sim.abundances <-round(rlnorm(5000, meanlog=2, sdlog=1))

#use this log-normal abundance curve to create a community

#data matrix (cdm) with 16 quadrats of species richness

#between 10 and 25.

cdm <-simulateComm(tree, min.rich=10, max.rich=25, abundances=sim.abundances)

#generate a phylogenetic distance matrix

dists <-cophenetic(tree)

#calculate the various forms of MPD using metricTester

naw.mpd <-modifiedMPD(cdm, dists, abundance.weighted=FALSE)

inter.mpd <-modifiedMPD (cdm, dists, abundance.weighted=“interspecific”)

intra.mpd <-modifiedMPD (cdm, dists, abundance.weighted=“intraspecific”)

#calculate the various forms of Clarke and Warwick’s

#metrics

temp.CW <-taxondive(cdm, dists) delta <-temp.CW$D

delta.star <-temp.CW$Dstar delta.plus <-temp.CW$Dplus

#Non-abundance-weighted MPD is equal to delta +. Also, call

#the raw values if you want to see those directly

plot(delta.plus∼naw.mpd)

#Interspecific abundance-weighted MPD is equal to delta *

plot(delta.star∼inter.mpd)

#Intraspecific abundance-weighted MPD is equal to delta

plot(delta∼intra.mpd)
~~~

**Figure S2.1.**
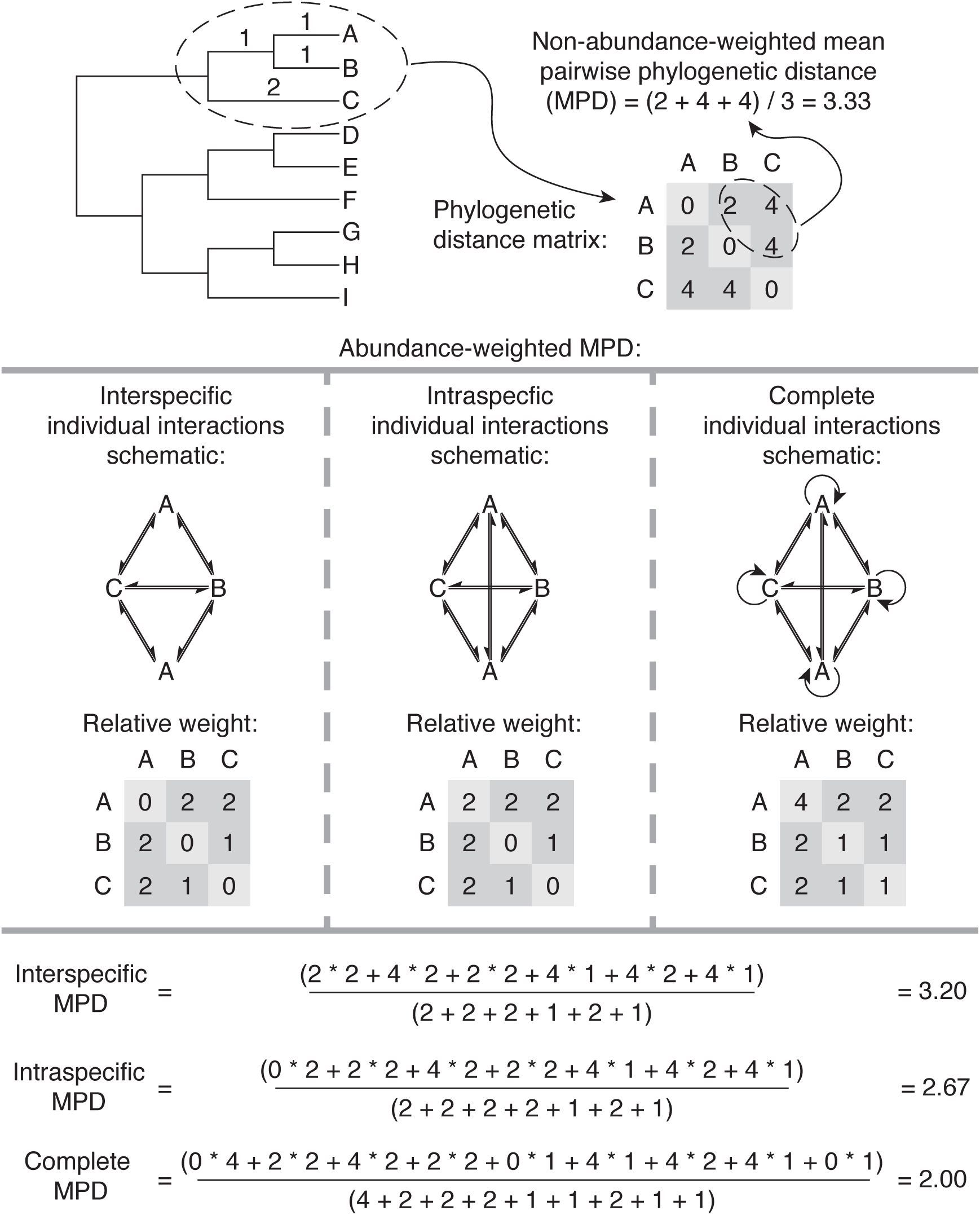
Schematic illustrating how non-abundance-weighted and three different forms of abundance-weighted MPD are calculated. Interspecific MPD accounts only for phylogenetic distances among heterospecifics, intraspecific also accounts for distances among conspecifics, and complete also includes interactions of an individual with itself.

**Figure S2.2.**
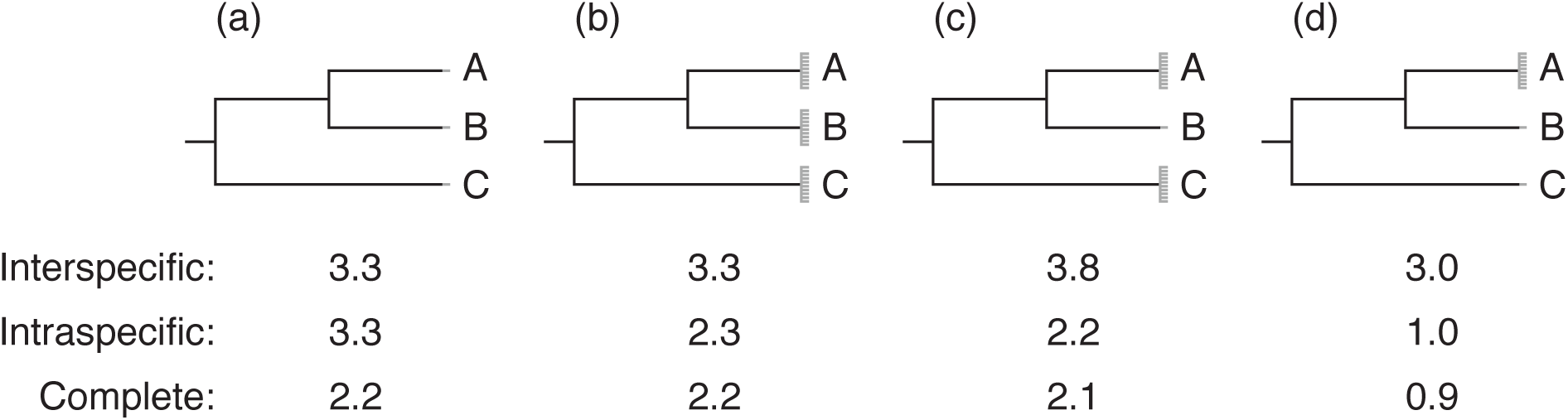
Examples showing how varying species’ abundances affects the different abundance-weighted MPD metrics. Branch lengths are the same as in Fig. S2.1. In all examples shown, unweighted MPD would be equal to 3.3. (a) Intraspecific MPD is equivalent to unweighted MPD in the special circumstance where one individual of each species is present, whereas the interspecific method is always equivalent to unweighted MPD when all species are equally abundant. (b) When all species’ abundances are increased, keeping relative abundances constant, intraspecifc MPD decreases as more intraspecific distances are incorporated. (c) When individuals are added to species A and C, interspecific MPD increases, emphasizing the distance between these upweighted species. Intraspecific and complete MPD decrease, emphasizing the intraspecific phylogenetic distances within species A and C. (d) Intraspecific and complete MPD decrease dramatically when only individuals of species A are added, whereas interspecific MPD decreases only somewhat (as a result of a down-weighting of the phylogenetic distance between species B and C).

**Figure S2.3.**
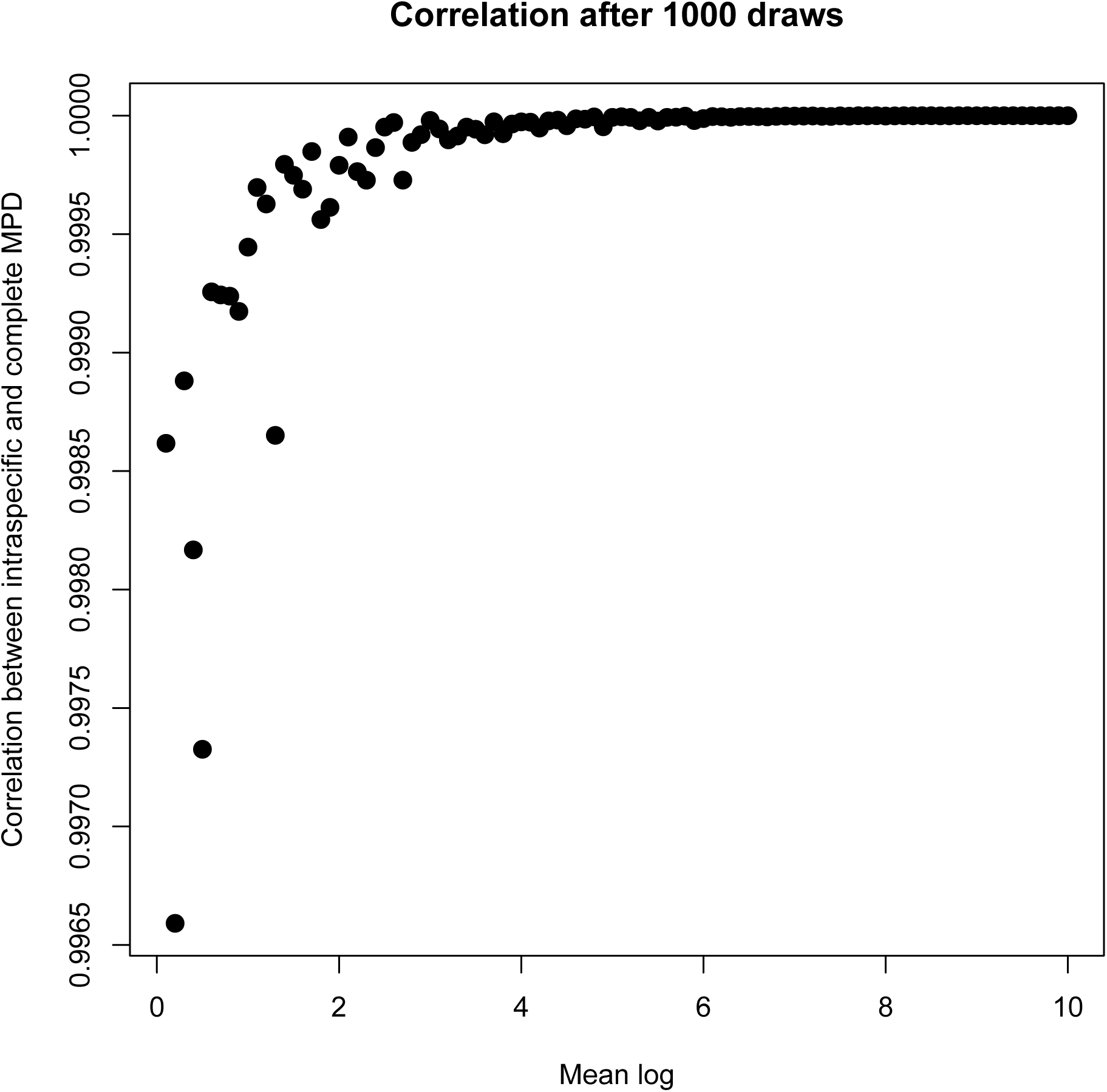
Intraspecific abundance-weighted MPD converges on complete abundance-weighted MPD with increasing total community size. To determine this, a series of 1,000 community data matrices were generated with the same phylogeny, number of species and number of quadrats, but the cells in the matrix were randomly filled by drawing from log-normal distributions with increasingly larger means.

**Figure S2.4.**
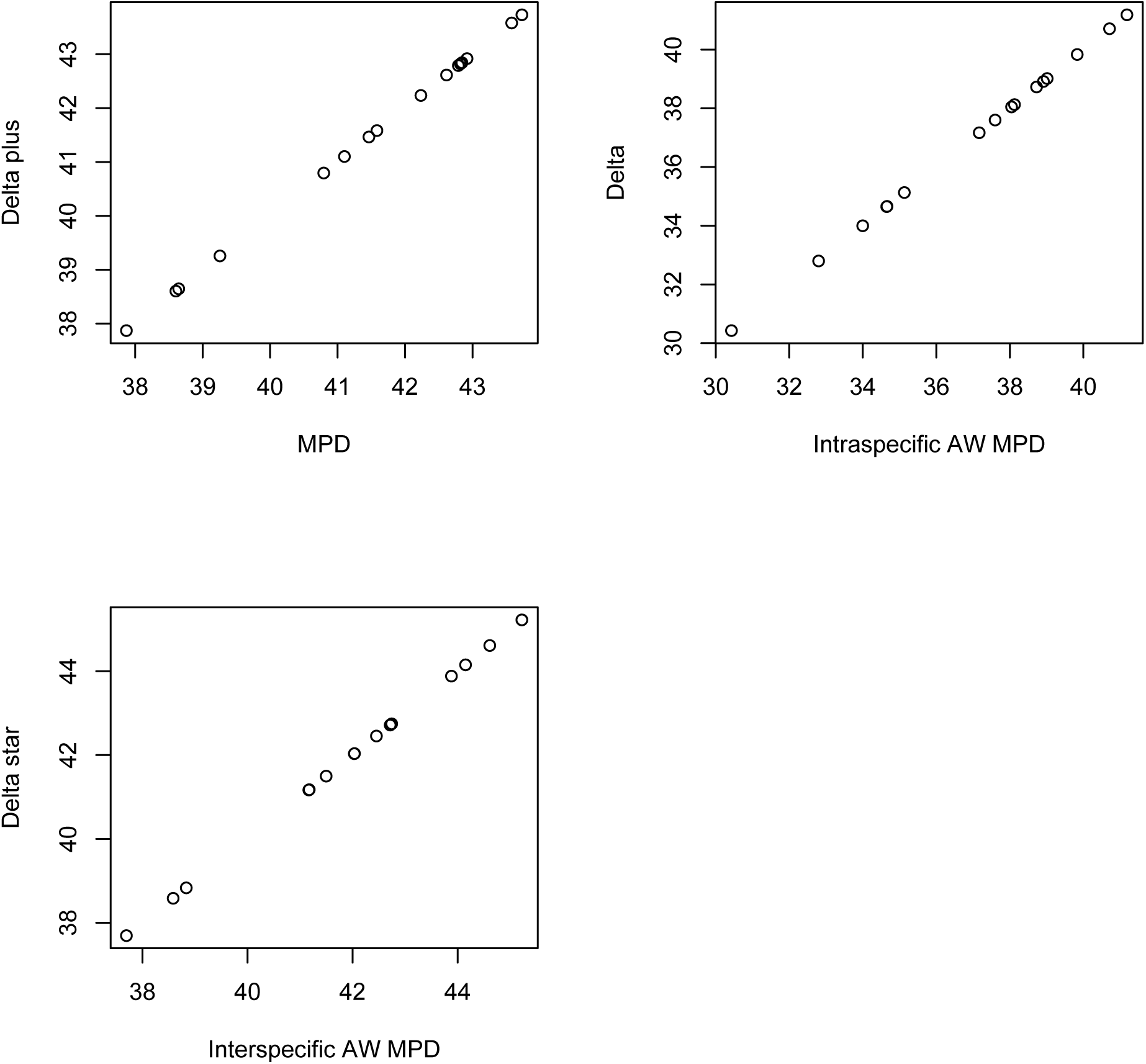
Scatterplots demonstrating the equivalency of Δ+ to MPD, Δ to intraspecific AW MPD, and Δ* to interspecific AW MPD. These plots were produced with the example code from *metricTester* shown above.

**1.**
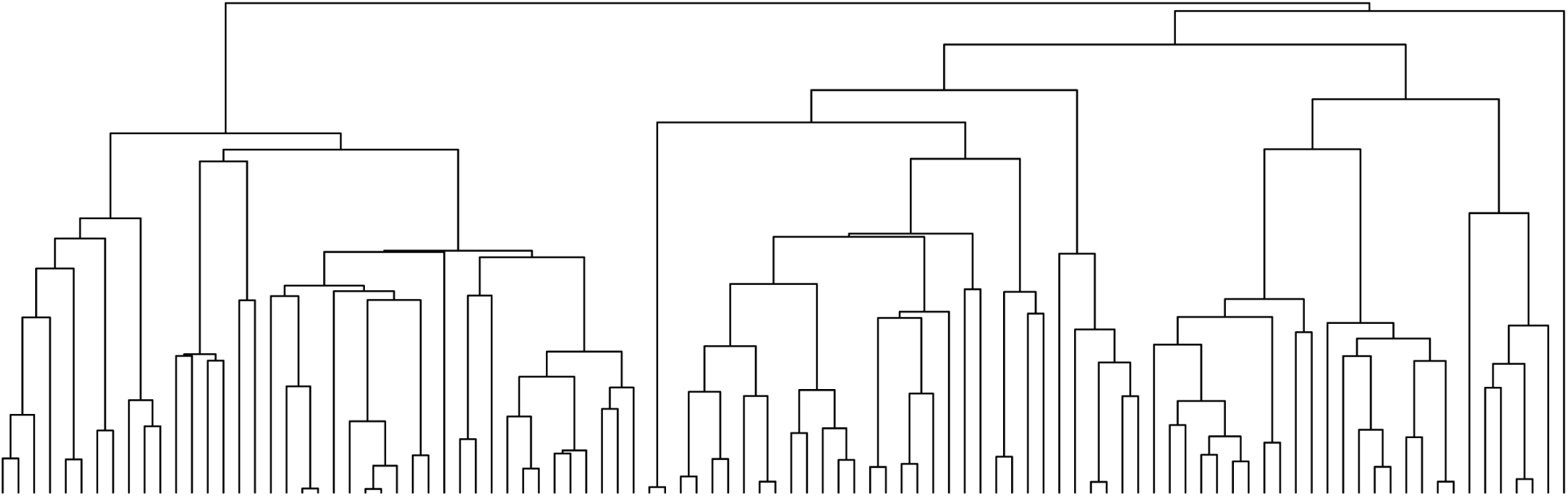
Generate phylogenetic tree

**2.**
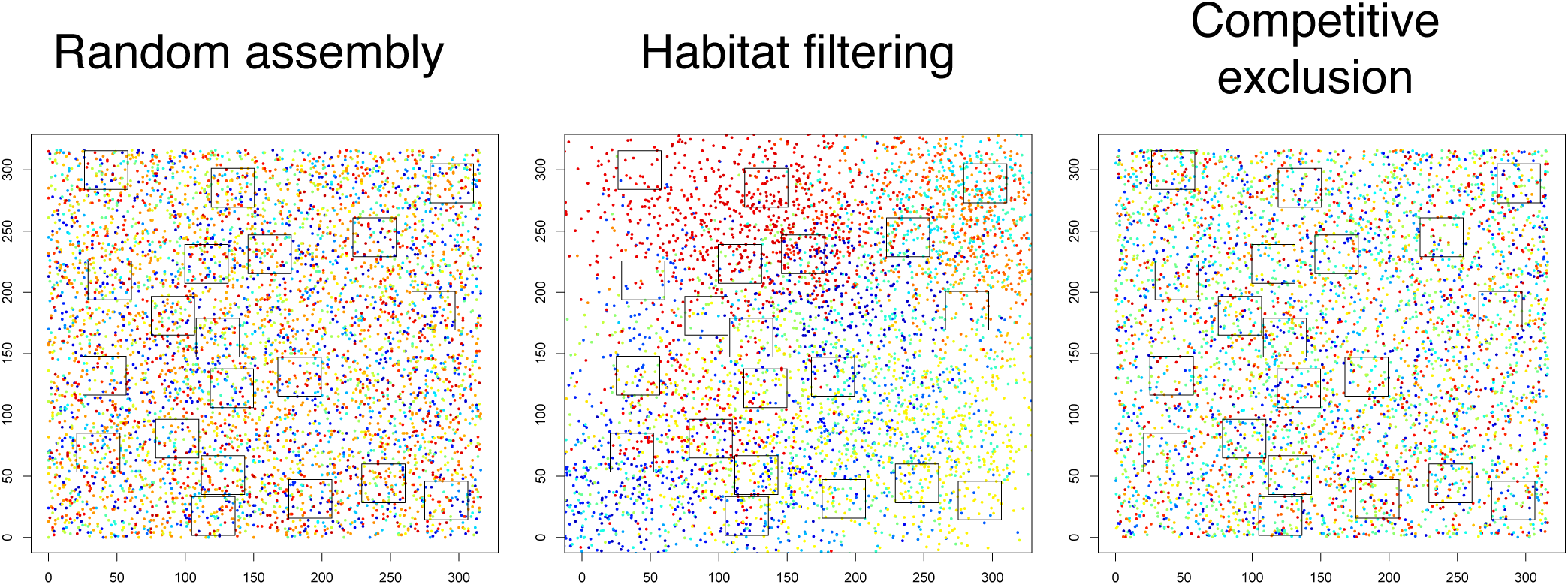
Run spatial simulations

Henceforth, schematic will only follow the random spatial assembly

**3.**
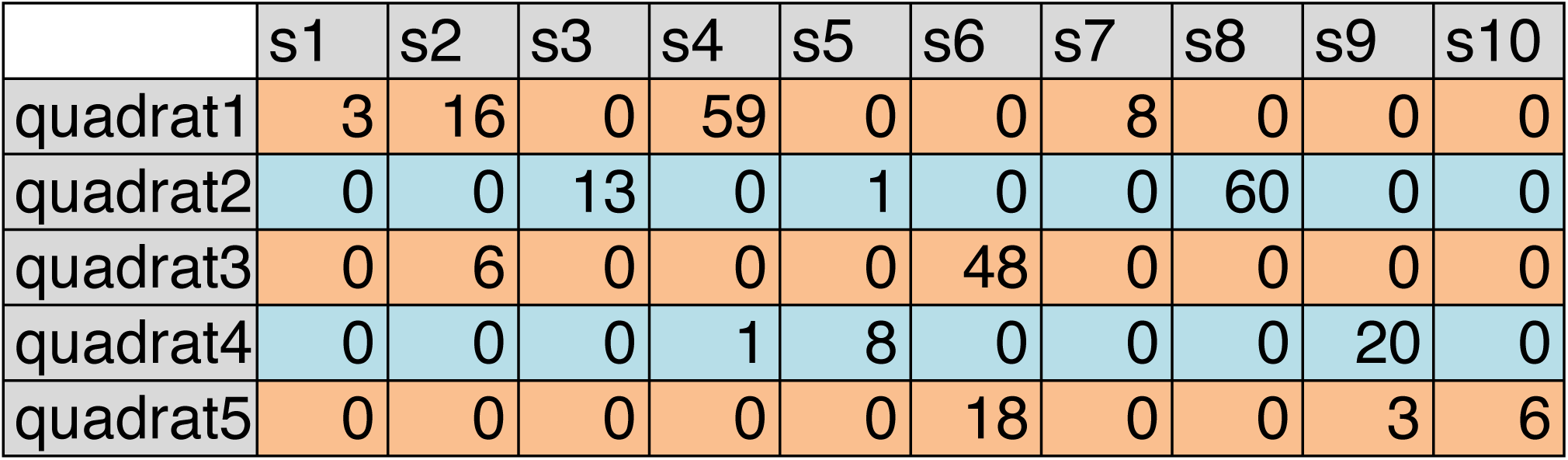
Sample quadrats and define CDM (real CDMs have 20 quadrats and up to 100 species)

**4.** Calculate observed metrics and set aside (only showing some of the metrics here)

| quadrat | richness | PSV | PSC | PSE | PAE | IAC | Haed | PD |
| --- | --- | --- | --- | --- | --- | --- | --- | --- |
| quadrat1 | 4 | 0.29 | 0.15 | 0.2 | 1.3 | -25 | 4.17 | 2.75 |
| quadrat2 | 3 | 0.31 | 0.28 | 0.16 | 0.94 | -42 | 3.56 | 2.63 |
| quadrat3 | 2 | 0.35 | 0.35 | 0.14 | 1 | -42 | 3.52 | 2.24 |
| quadrat4 | 3 | 0.19 | 0.13 | 0.16 | 1.4 | -6.3 | 3.14 | 2.18 |
| quadrat5 | 3 | 0.75 | 0.49 | 0.57 | 0.84 | -11 | 3.14 | 3.71 |

**5.**
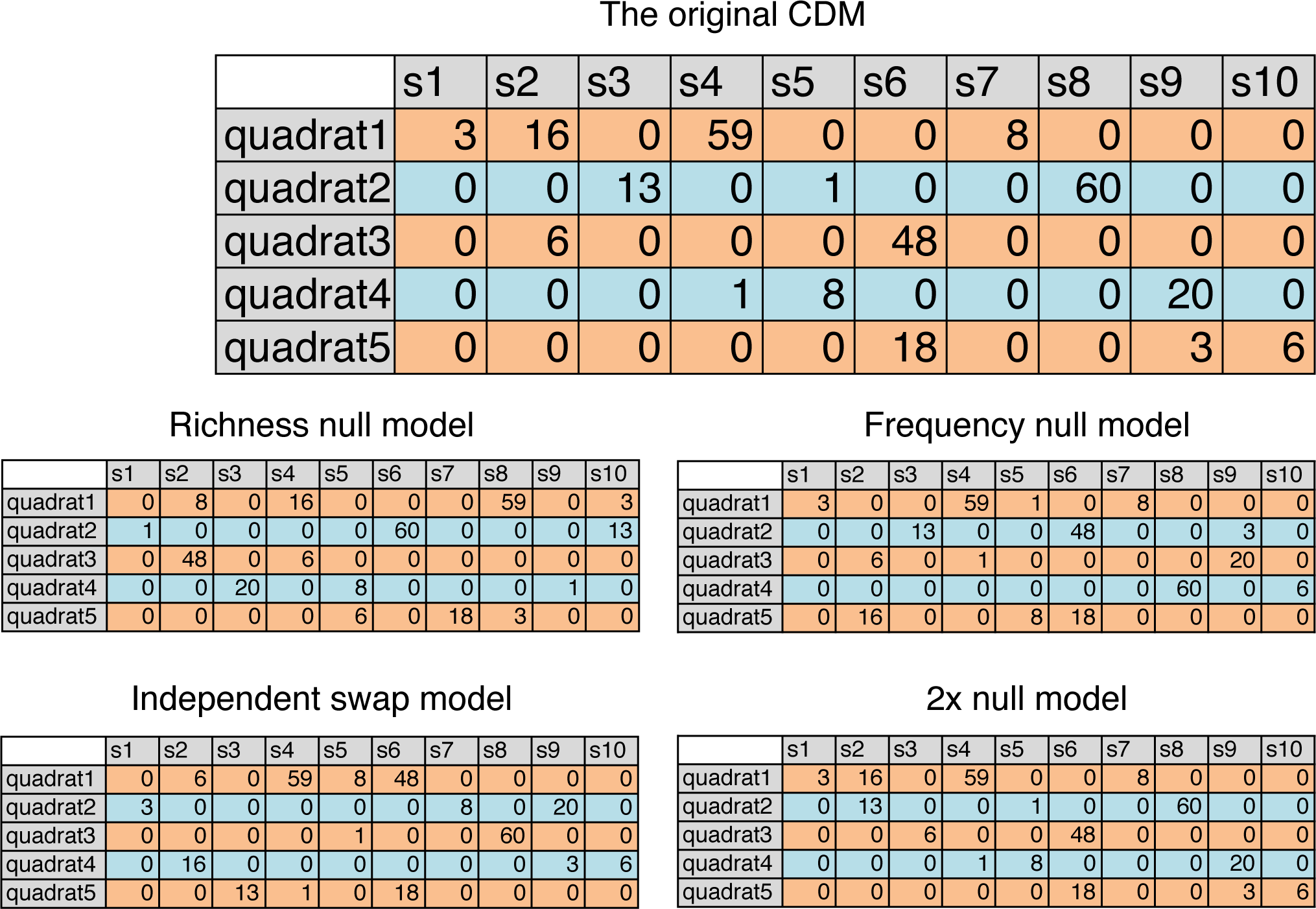
Shuffle original CDM according to each null model (only showing some of the null models here). Use these matrices to recalculate the metrics.

Henceforth, schematic will only follow the richness null model

**6.**
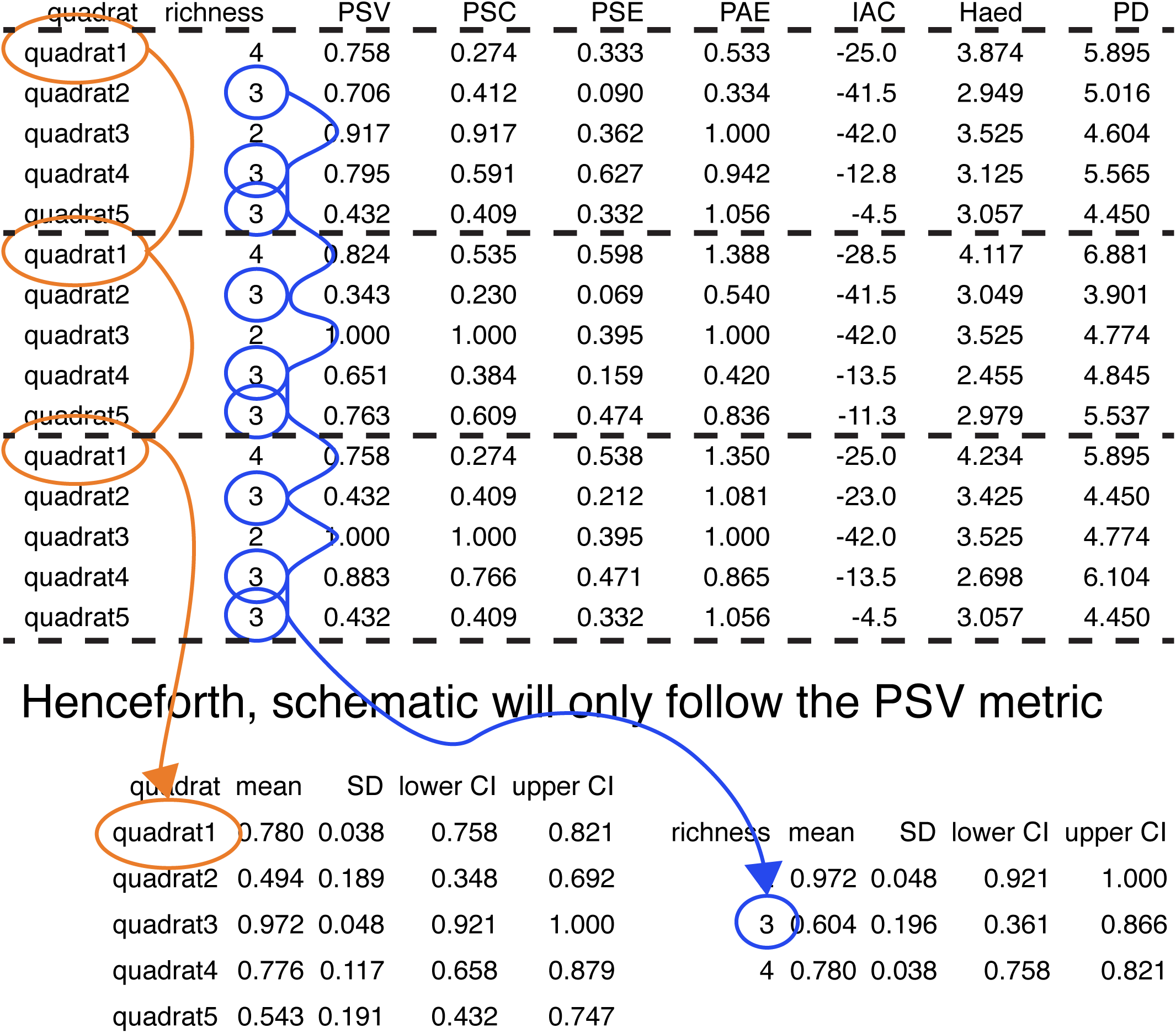
Repeat CDM randomizations 1,000 times, calculating all metrics each randomization, and retain all randomized values (i.e. append to a growing data frame). Three randomizations are shown here. 7. Concatenate and summarize randomized values by both quadrat and by richness

Henceforth, schematic will only follow concatenation by quadrat method

**8.**
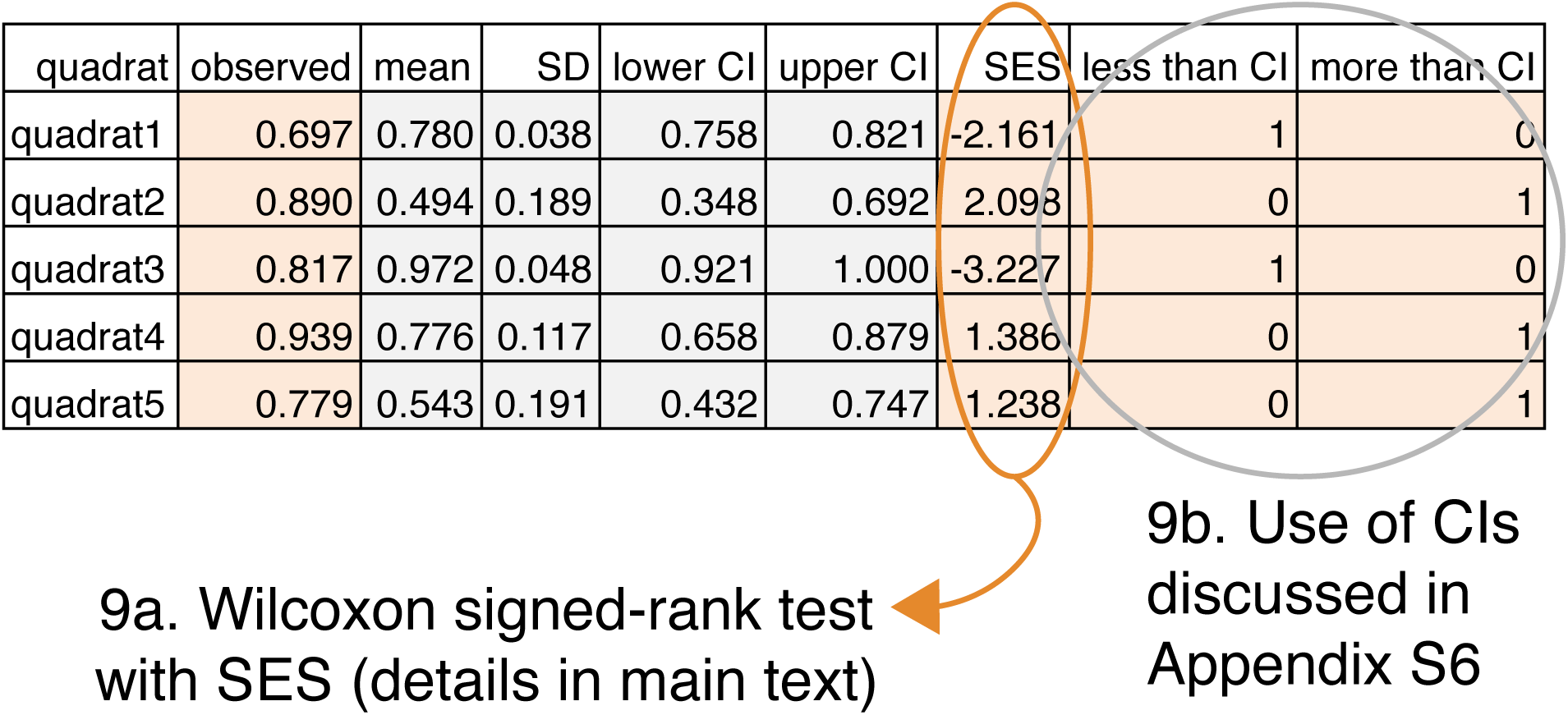
Use these summaries to generate SES and determine whether a given quadrat deviates beyond CIs

This diagram only followed the random spatial simulation, the richness null model and the PSV metric concatenating by quadrat method from start to finish. It also only followed 5 quadrats. In practice, a full, single iteration tests 20 quadrats and employs the random, filtering and competition simulations, all 9 null models, all 22 metrics, and concatenates by both richness and quadrat, then calculates SES and CIs, and uses these to assess significance. We ran 1,009 such iterations.

## Appendix S4

Metric and null model approach results are robust to variation in spatial simulation parameters and number of randomizations of community data matrix.

In this study we simulated community assembly and derived a community data matrix (CDM) from the simulated spatial arena. We simulated these communities according to one of three assembly processes: random assembly, habitat filtering and competitive exclusion. Observed phylogenetic community structure metrics from the CDM were then compared to null distributions of the metrics generated by randomizing the observed CDM 10^3^ times. Results were very similar across metrics and nulls when the number of randomizations of each CDM was increased from 10^3^ to 10^4^, with slight shifts in error rates for some approaches (Fig. S4.1). That said, we do encourage empirical researchers to use more than 10^3^ randomizations of their observed CDM, e.g., 10^6^.

Communities in the main results contained 2,000-4,000 individuals (i.e. 200-400/ha). This is around a quarter to a half of the number of individual stems recorded in Australian forest plots of similar spatial extent (Murphy et al. 2013), and much less than the 6,400 individuals/ha in Ecuadorian rainforests (Valencia et al. 2004). However, our results were robust to increasing the number of individuals placed per community to between 7,000 and 11,000 (Fig. S4.2).

The competitive exclusion simulations were robust to parameter variation.

Variation in both the interaction distance and in the percent of individuals “killed” per generation resulted in communities with regions of approximately equal genetic “overdispersion” (Fig. S4.3-4). These communities looked similar to the random communities, though the even spacing of close relatives was discernible (Fig. S4.5).

During the competitive exclusion simulations, some species that were initially common in the community became less so with each generation (Fig. S4.6). These species were those with many close relatives in the phylogeny. A null model like the independent swap that incorporates species occurrence frequencies derives these from occurrence frequencies in the observed community data matrix (CDM). After the competitive exclusion simulations, therefore, longer than average branch lengths end up being frequently sampled in the randomized CDMs. Accordingly, the expected phylogenetic community structure is shifted upwards from that given a richness null, and it becomes difficult to detect phylogenetic overdispersion (Fig. S1.7). This occurs despite the fact that, throughout the competitive exclusion simulations, removed individuals are settled from the initial regional abundance pool. Our development of the regional null model (Appendix S1) was motivated in large part by this complication.

R code for example communities simulated with *metricTester* (and the specific parameters we used in the main results) is given below:

Simulate a phylogeny of 100 species with geiger:

~~~
tree <-sim.bdtree(b=0.1, d=0, stop=“taxa”, n=100)
~~~

Generate an object of class “simulations.input”:

~~~
prepped <-prepSimulations(tree=tree,
arena.length=sqrt(100000), mean.log.individuals=3.5,
length.parameter=1000, sd.parameter=40, max.distance=20,
proportion.killed=0.2, competition.iterations=60)
~~~

Run whatever simulations have been defined in the function defineSimulations(). In our case, this would simultaneously run the random, habitat filtering and competitive exclusion simulations, but more can be easily defined and added by users. Examples of what these simulated communities look like can be seen in Appendix S3.

~~~
simulations <-runSimulations(prepped)
~~~

**Figure S4.1.**
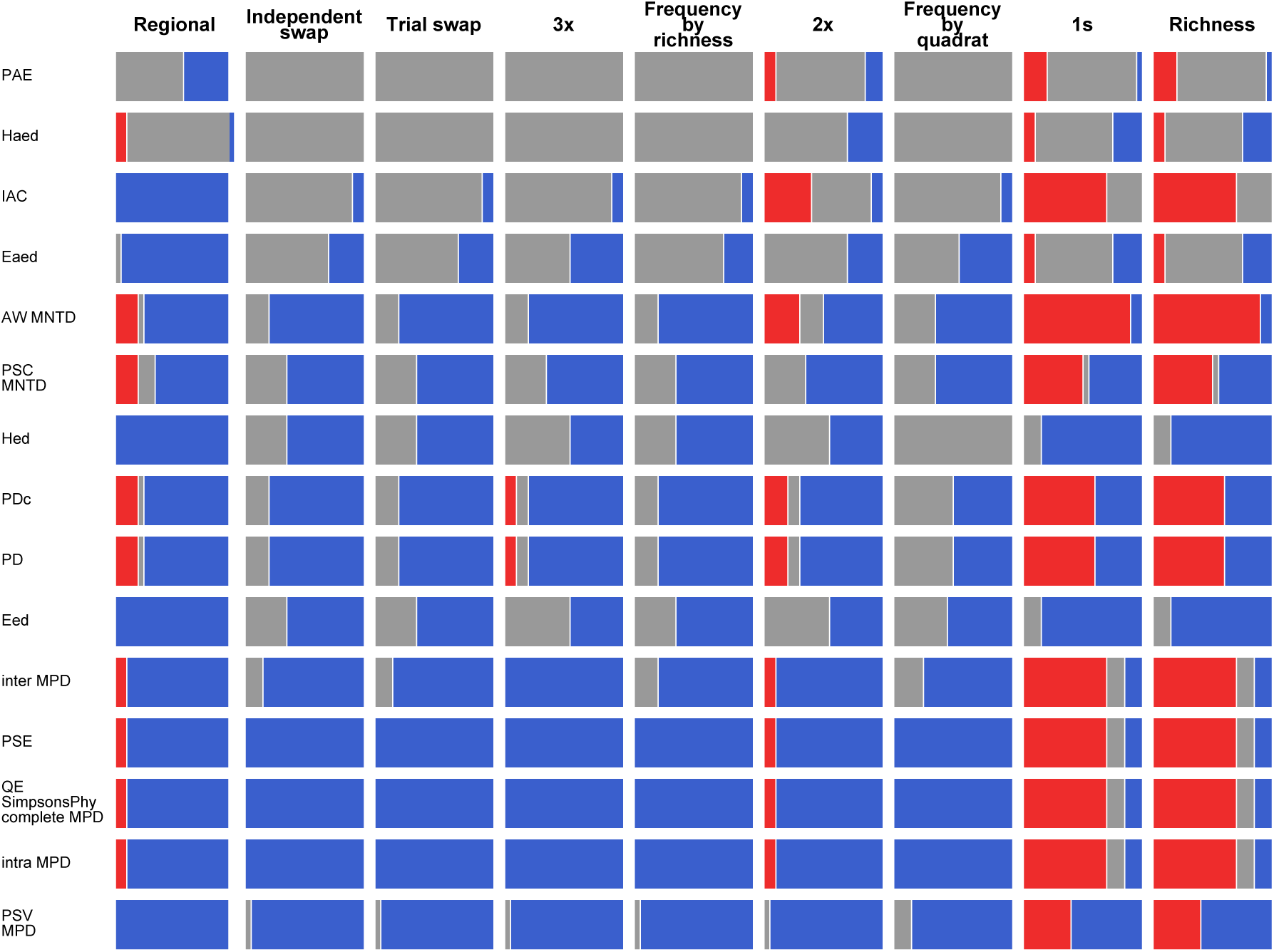
Overall performance of metric + null model approaches given spatial simulations where the resulting CDM was randomized 10^4^ instead of 10^3^ times. This figure is analogous to Fig. 5 in main text. Red bars (type I errors) summarize the proportion of 10 total random community assembly simulations where the mean of the standardized effect sizes differed significantly from zero (two-way Wilcoxon signed-rank test). Gray bars summarize the mean type II error rates from the 10 habitat filtering and competitive exclusion simulations. Blue bars provide an indication of the success of each approach, and are defined as the proportion of runs that did not generate type I and II errors. Metrics and nulls are arranged in order from overall best-performing to worst, with the best approaches in the bottom left corner. Results are generally very similar to those in the main text, though there are notable improvements in the performance of *E*_ED_ and *H*_ED_ and declines in the performance of PAE and PD.

**Figure S4.2.**
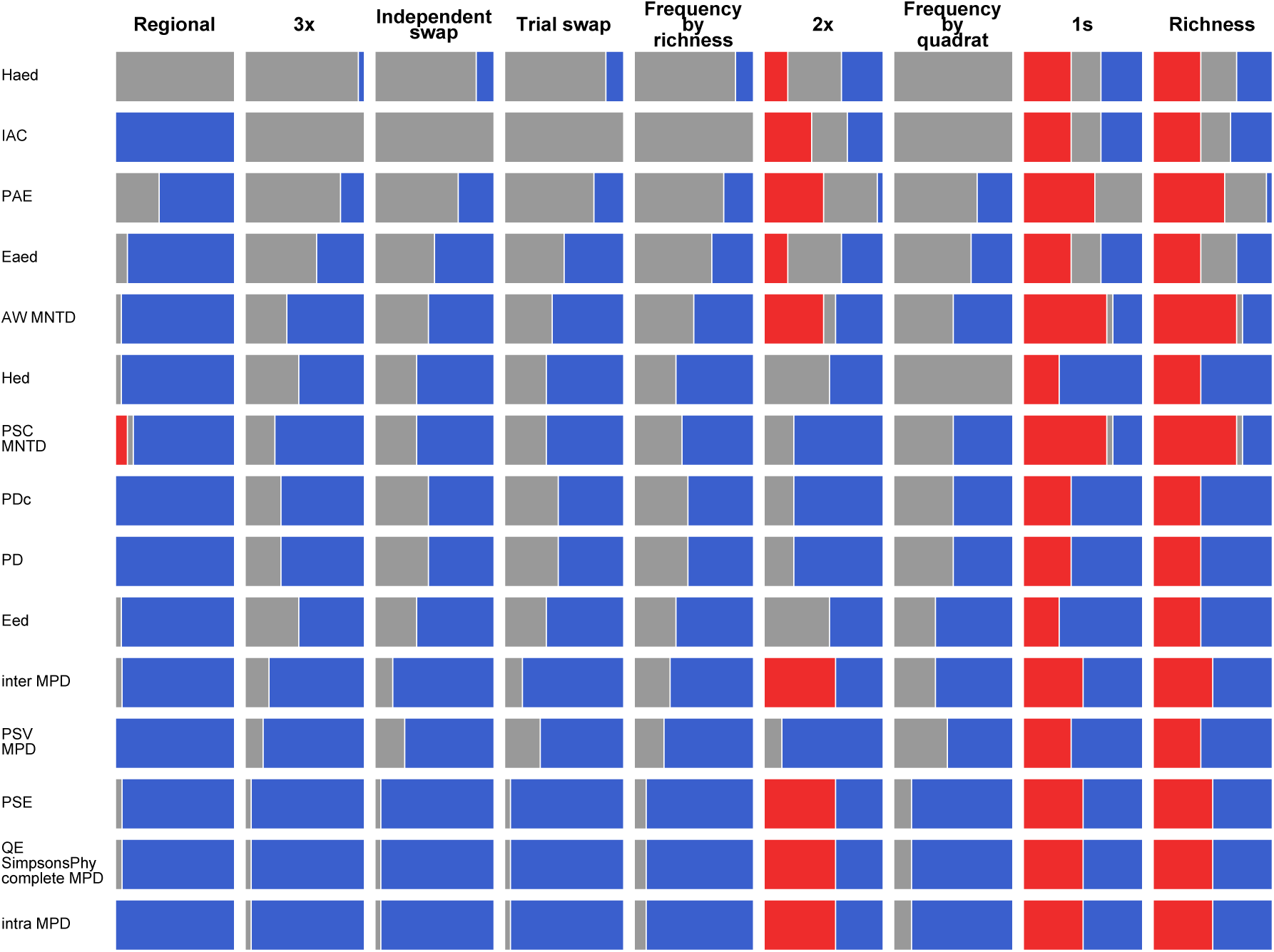
Overall performance of metric + null model approaches given spatial simulations containing more individuals (8,000 – 11,000 instead of 2,000 – 4,000), analogous to Fig. 5 in main text. As compared with the example code in this appendix, these simulations are generated by changing the mean.log.individuals argument to 4. Red bars (type I errors) summarize the proportion of 10 total random community assembly simulations where the mean of the standardized effect sizes differed significantly from zero (two-way Wilcoxon signed-rank test). Gray bars summarize the mean type II error rates from the 10 habitat filtering and competitive exclusion simulations. Blue bars provide an indication of the success of each approach, and are defined as the proportion of runs that did not generate type I and II errors. Metrics and nulls are arranged in order from overall best-performing to worst, with the best approaches in the bottom left corner. Results are generally very similar to those in the main text, though there are notable improvements in the performance of *E*_ED_ and *H*_ED_ and declines in the performance of the 2x null model.

**Figure S4.3.**
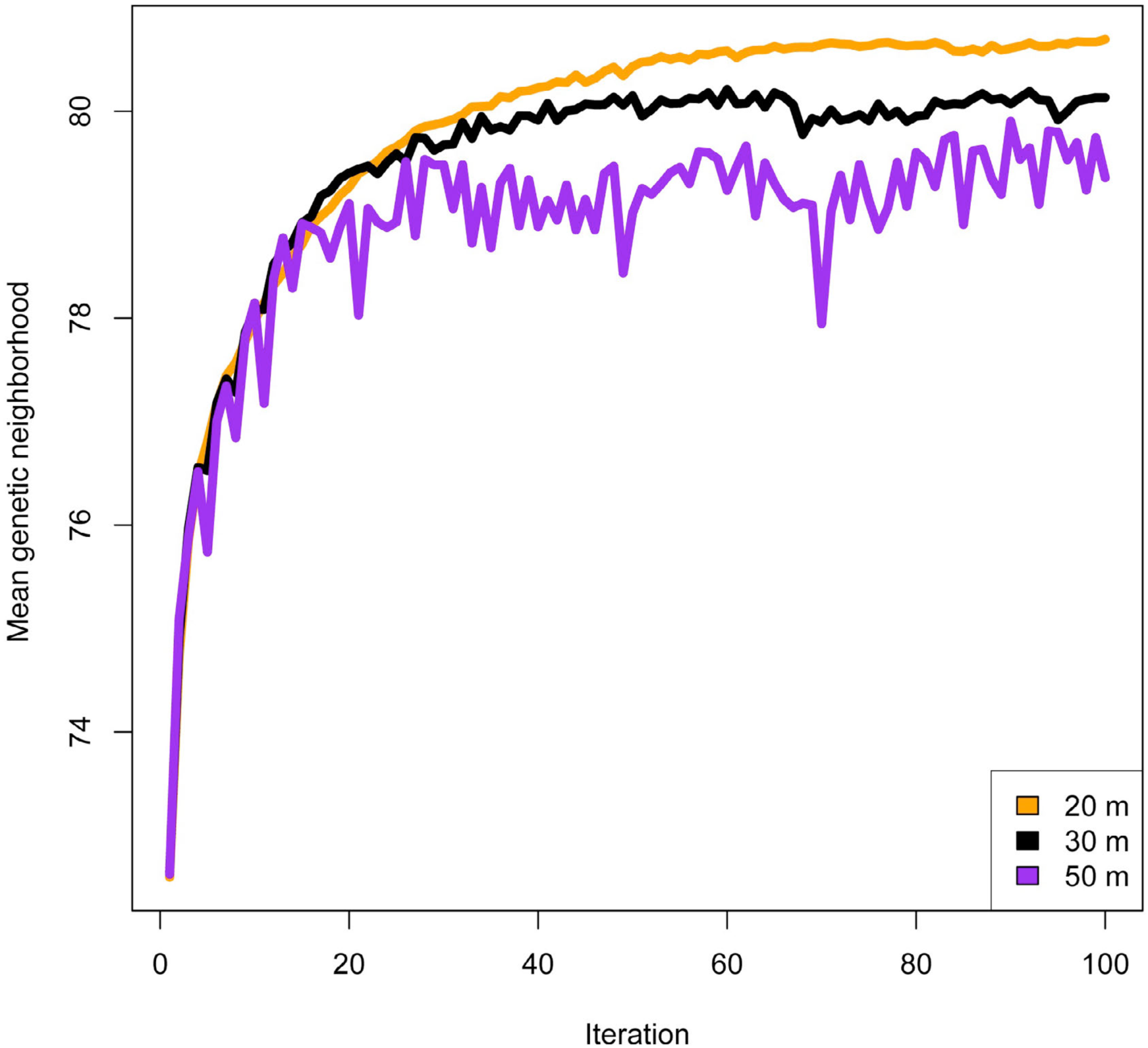
Change in the mean genetic neighborhood over 100 generations of the competitive exclusion assembly for three different interaction distances. For each individual in the community, the mean pairwise phylogenetic distance between itself and all individuals within the interaction distance is derived. The mean genetic neighborhood is then defined as the community mean of these individual means. Across a wide range of interaction distances, the general pattern of increasing phylogenetic overdispersion is evident.

**Figure S4.4.**
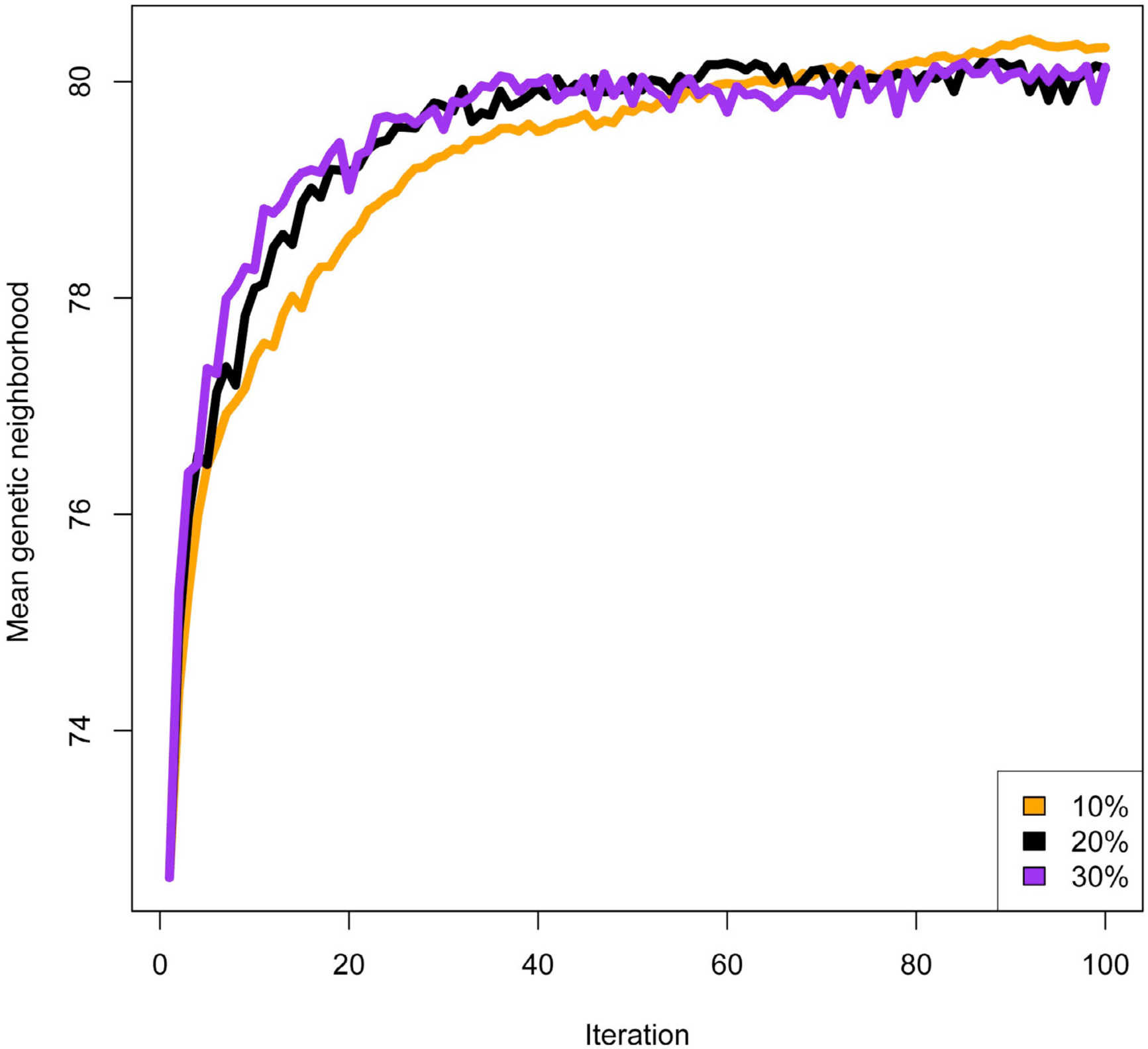
Change in the mean genetic neighborhood over 100 generations of the competitive exclusion assembly for three different interaction distances. For each individual in the community, the mean pairwise phylogenetic distance between itself and all individuals within the interaction distance is derived. The mean genetic neighborhood is then defined as the community mean of these individual means. Across a wide range of percent killed parameters, the general pattern of increasing phylogenetic overdispersion is evident. Based on these preliminary results, it appears that removing (“killing”) a small percentage (e.g., 10%) of individuals each generation would ultimately generate a similar pattern to removing a large percentage (e.g., 30%).

**Figure S4.5.**
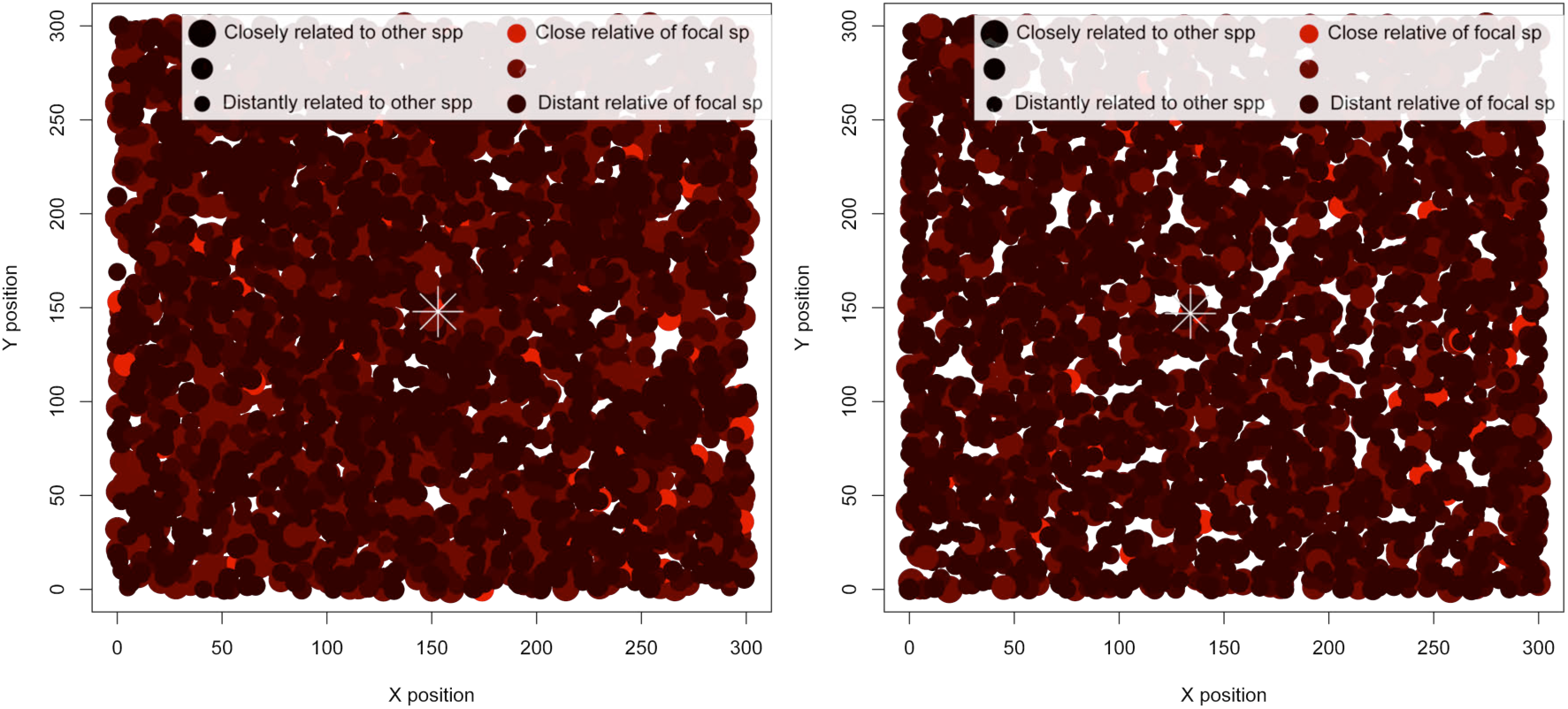
On the left, an example of a 300 × 300 m random assembly community, created using similar parameters to those in the study. Here, a random individual was selected near the center of the community (marked with a white asterisk). Individuals were then color-coded as a function of their relatedness to the focal individual, where bright red indicates a member of the same species. The size of individual dots was scaled according to their mean relatedness to all other species in the phylogeny, such that large dots indicate a member of a species with many close relatives. In this random community, bright red dots occasionally occur close together, and on average the plot is “redder” than the right panel. Also, the dots in the plot appear to be more uniform in size. On the right, an example of a 300 × 300 m competitive exclusion community. The community from the left panel was used as a starting point. An individual of the same focal species as that panel was selected near the center of the community (marked with a white asterisk). Size and color scaling are the same as in the left panel. In this community, bright red dots appear regularly spaced, and on average the plot is “darker” than the left panel. Also, the dots in the plot appear to be more heterogeneous in size.

**Figure S4.6.**
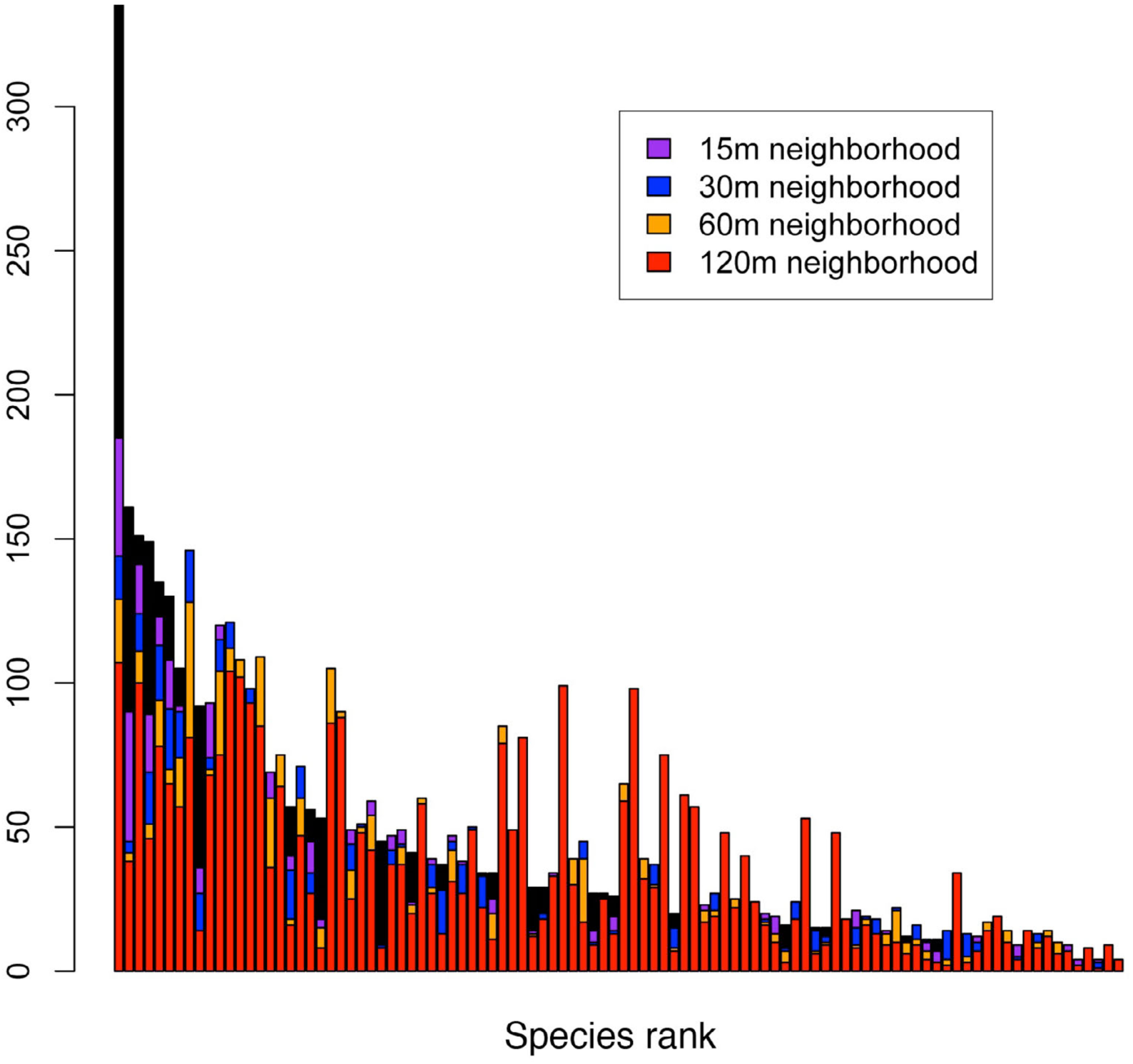
Changes in the rank abundance curve after 25 generations of the competitive exclusion assembly simulations. The initial rank abundance curve is shown in black. Increasing the interaction distance results in increasingly large deviations from the initial rank abundance curve. Some species (e.g., columns 8 and 9) change abundance dramatically during these competition simulations. Four separate simulations with the same initial community and phylogeny are shown here.

**Table S5.1.**
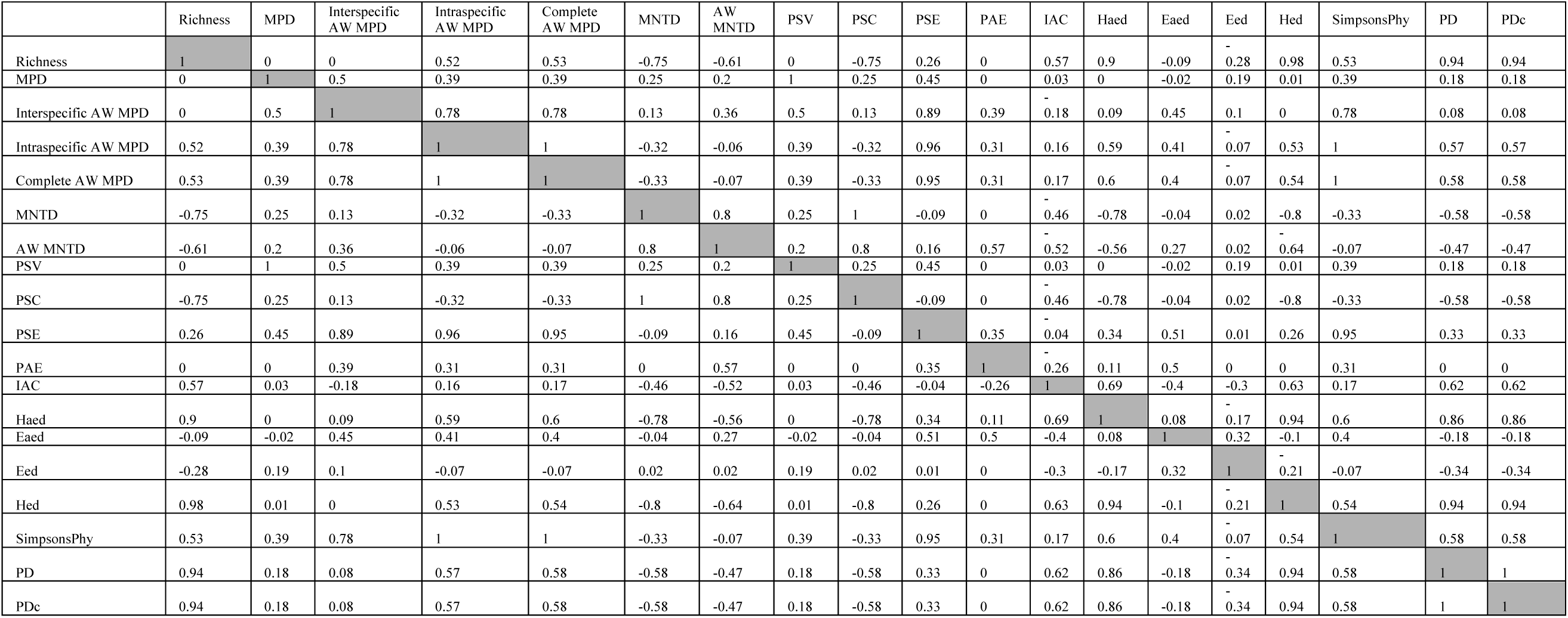
Pearson correlation coefficients among 19 phylogenetic community structure metrics. The raw values used to calculate these correlations were generated over 50,000 randomized community data matrices spanning a range of species richness values from 10 to 40.

## Appendix S6

Expanded methods and results for approaches that concatenate randomized values by richness and approaches that assess significance on a per-quadrat basis.

As in the main text, we here define a community data matrix (CDM) as a quadrat (rows)-by-species (columns) matrix, where cells are filled according to the abundance of a given species in a given quadrat. Traditionally, with null models used in empirical studies of phylogenetic community structure, the CDM is randomized according to some algorithm (e.g., the independent swap), and the metric in question is recalculated row-wise after each randomization. These randomizations are performed some large number of times, the quadrat-specific expectations are then compared to observed values, used to derive standardized effect sizes (SES), and significance of the observed matrix-wide deviation of SES from expectations is assessed with a test such as a Wilcoxon signed-rank test.

A slight deviation from this approach is to assess significance on a per-quadrat basis (Miller et al. 2013). Here, a given quadrat is considered significantly overdispersed or clustered if it deviates above or below, respectively, the 95% confidence intervals (CI) for that quadrat based on the randomized values from the null model. An additional deviation from the traditional approach is to retain randomized values, along with the species richness (the row-wise sum of non-zero elements) of the corresponding quadrat from the randomized matrix, and concatenate and summarize randomized metric values by these species richness values.

Thus, null models such as the richness, independent swap and trial swap (Table 2, main text) that maintain row-wise sums of non-zero elements are expected to perform similarly whether results are concatenated by richness or quadrat. Indeed, since a given species richness may occur more than once in an observed CDM (e.g., two sampled quadrats from a given community might both contain 20 species), running a null model in this manner can effectively increase the number of randomizations against which observed values are compared. In other words, the expectations for a given observed quadrat of 20 species should be no different than those from a different quadrat of 20 species. However, other null models like the frequency null, which do not maintain row-wise sums of non-zero elements, are expected to behave differently when concatenated by richness or quadrat. This was the impetus for the development of the frequency by richness null (Miller et al. 2013), which we showed here to be equivalent to the independent and trial swap null models.

During our simulations, we performed all analyses according both to a traditional SES, community-wide framework, and to a per-quadrat significance framework. We also performed both of these types of analyses after concatenating randomized metric values both by quadrat and by species richness. Thus, in addition to the overall results presented in the main text (Fig. 5), there were three other ways to consider the overall results: (1) a SES framework where results were concatenated by species richness, (2) a per-quadrat framework where results were concatenated by quadrat, and (3) a per-quadrat framework where results were concatenated by species richness.

For the latter two approaches, we summarized type I errors for a given metric + null approach as the sum of all clustered or overdispersed quadrats from the random simulation, overdispersed quadrats from the filtering simulation, and clustered quadrats from the competitive exclusion simulation divided by the total number of quadrats from each of these three simulations. We summarized type II errors for a given metric + null approach as the sum of all not clustered quadrats from the filtering simulation and all not overdispersed quadrats from the competitive exclusion simulation divided by the total number of quadrats from both of these simulations.

While the traditional approach for calculating the significance of observed phylogenetic community structure is sufficient, there are at least three reasons researchers might choose to take one of the other approaches. First, a researcher might wish to use unstandardized metrics (e.g., MPD instead of NRI). Second, a researcher might wish to assess significance for a given quadrat instead of across an entire matrix. Of course, using a SES approach, a single quadrat with an SES of > |1.96| can be considered significant at an alpha of 0.05. However, if the researcher wishes to use unstandardized metrics, assessing significance as per-quadrat deviations beyond CI is a more appropriate approach. Third, we believe it is not well appreciated that null model expectations vary in metric-specific manners across species richness, and significance testing in this manner may provide insight into how such underlying expectations vary in the empirical dataset in question.

When we took an SES framework where results were concatenated by species richness, as expected, results were similar to those when concatenated by quadrat (Fig. 5, main text) for the trial swap, independent swap, richness and 1s null models (Fig. S6.1). They were identical for the regional and frequency by richness null models since these were also concatenated by richness in the main text, and they were identical for the frequency by quadrat null model since we only concatenated those randomized values by quadrat (Fig. S6.1). Unexpectedly, however, since both null models maintain species richness (i.e. row-wise sums of non-zero elements), results differed dramatically for both the 2x and 3x when randomizations were concatenated by richness.

We investigated this surprising result by creating a CDM of 12 quadrats and 20 species total, where quadrats either contained 5, 7, or 10 species each (4 quadrats of each richness). When we randomized the matrix according to the 2x and 3x null models and plotted per-quadrat expectations as a function of the species richness of that quadrat, we discovered that quadrats of similar species richness did not appear to converge on similar expectations. Thus, when randomizations from a given species richness were concatenated, instead of being (at least somewhat) normally distributed, the randomized values often exhibited notably multi-modal distributions (e.g., Fig. S6.2). Moreover, even randomized values from a given quadrat were sometimes multi-modal (e.g., Fig. S6.3). This explains the significant drop in power of the 2x and 3x null models when concatenated by richness—expectations at a given species richness tend to be more platykurtic than when concatenated by quadrat.

The drop in power of the 2x and 3x null models continued when the significance of observed metrics was assessed on a per-quadrat basis. With this manner of significance testing, the type II error rates of both null models was greater than 99.9% when randomizations were concatenated either by quadrat (Fig. S6.4) or by richness (Fig. S6.5).

Both the 2x and 3x null models performed well when used in a traditional manner. However, when results were either concatenated by richness and/or significance was assessed on a per-quadrat basis, these models essentially lacked any power to detect simulated community assembly processes. More study of the behavior of these models is warranted. Qualitatively, the issue appears to be biased exploration of plausible phylogenetic community structure space, possibly due to highly constrained randomization algorithms that maintain numerous aspects of the initial CDM.

**Figure S6.1.**
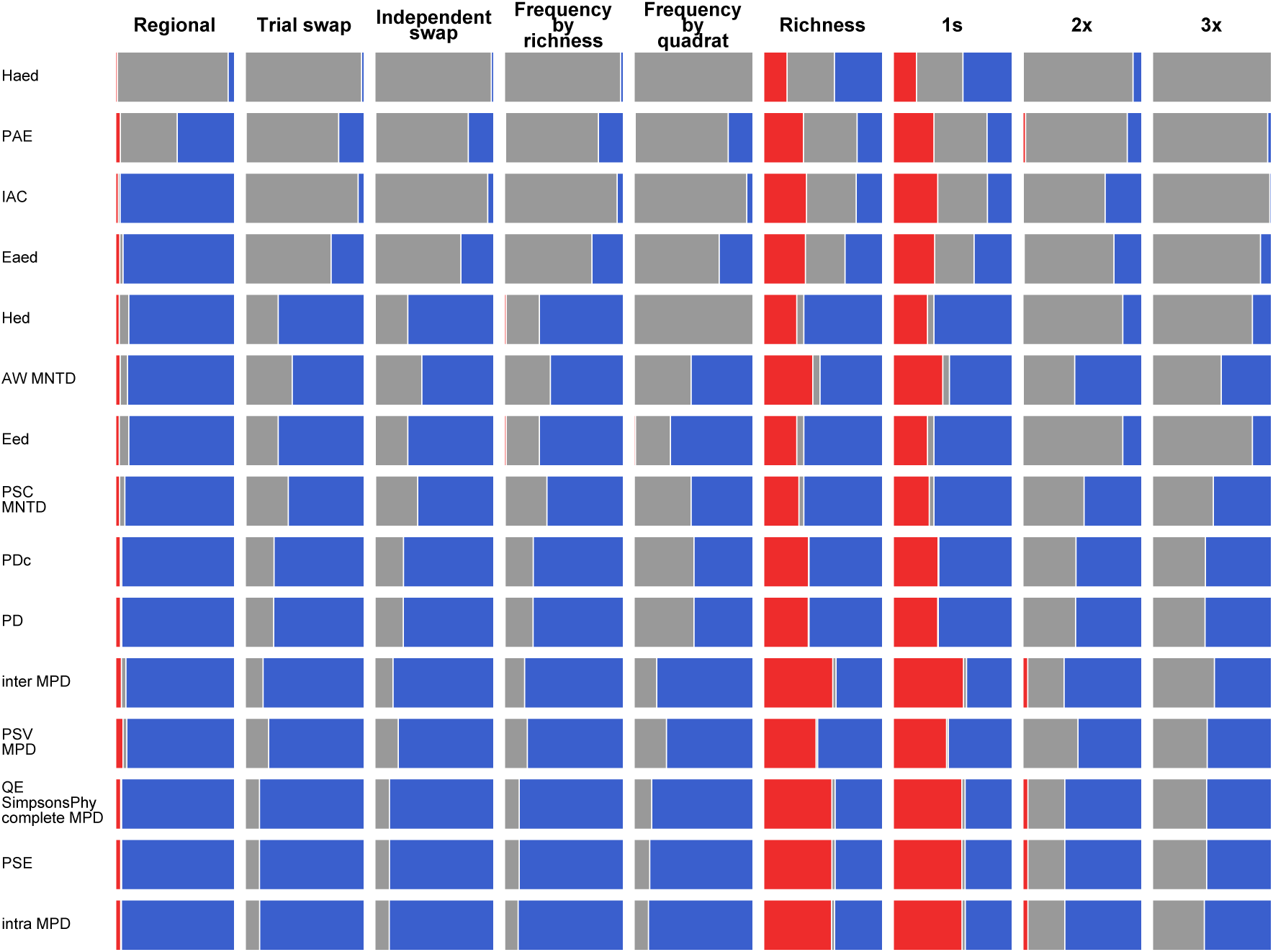
Overall performance of metric + null model approaches when randomizations were concatenated by richness (except for the frequency by quadrat model, which is always concatenated by quadrat). Red bars (type I errors) summarize the proportion of the total 1,009 random community assembly simulations where the mean of the standardized effect sizes differed significantly from zero (two-way Wilcoxon signed-rank test). Gray bars summarize the mean type II error rates from the habitat filtering and competitive exclusion simulations. Blue bars provide an indication of the success of each approach, and are defined as one minus the mean type I and II error rates. Metrics and nulls are arranged in order from overall best-performing to worst, with the best approaches in the bottom left corner.

**Figure S6.2.**
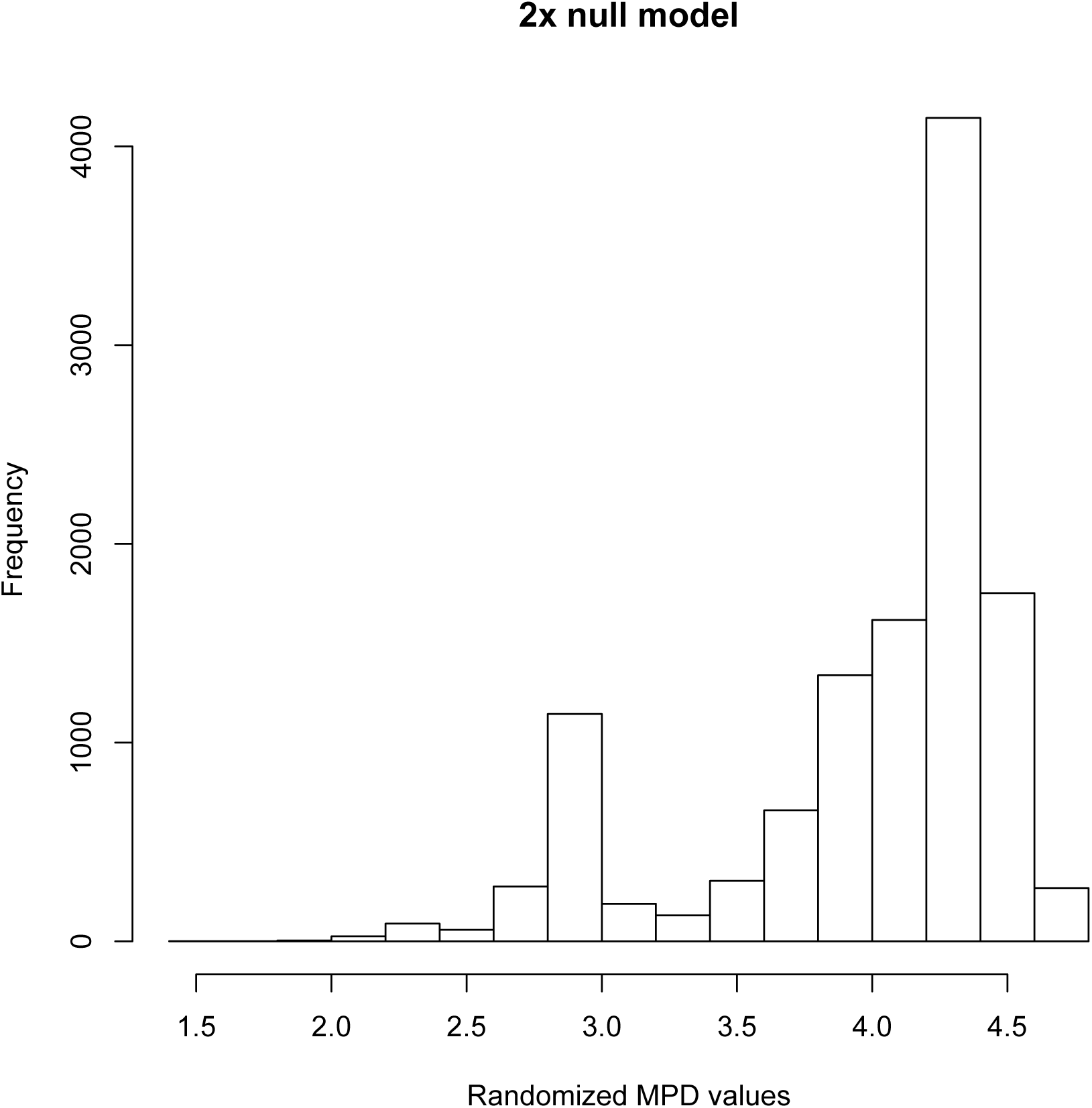
Distribution of randomized MPD values after 3,000 randomizations of the CDM described in this appendix. Here, all randomized values from the three quadrats that contained five species each are shown (i.e. there are 15,000 randomized MPD values shown in the histogram).

**Figure S6.3.**
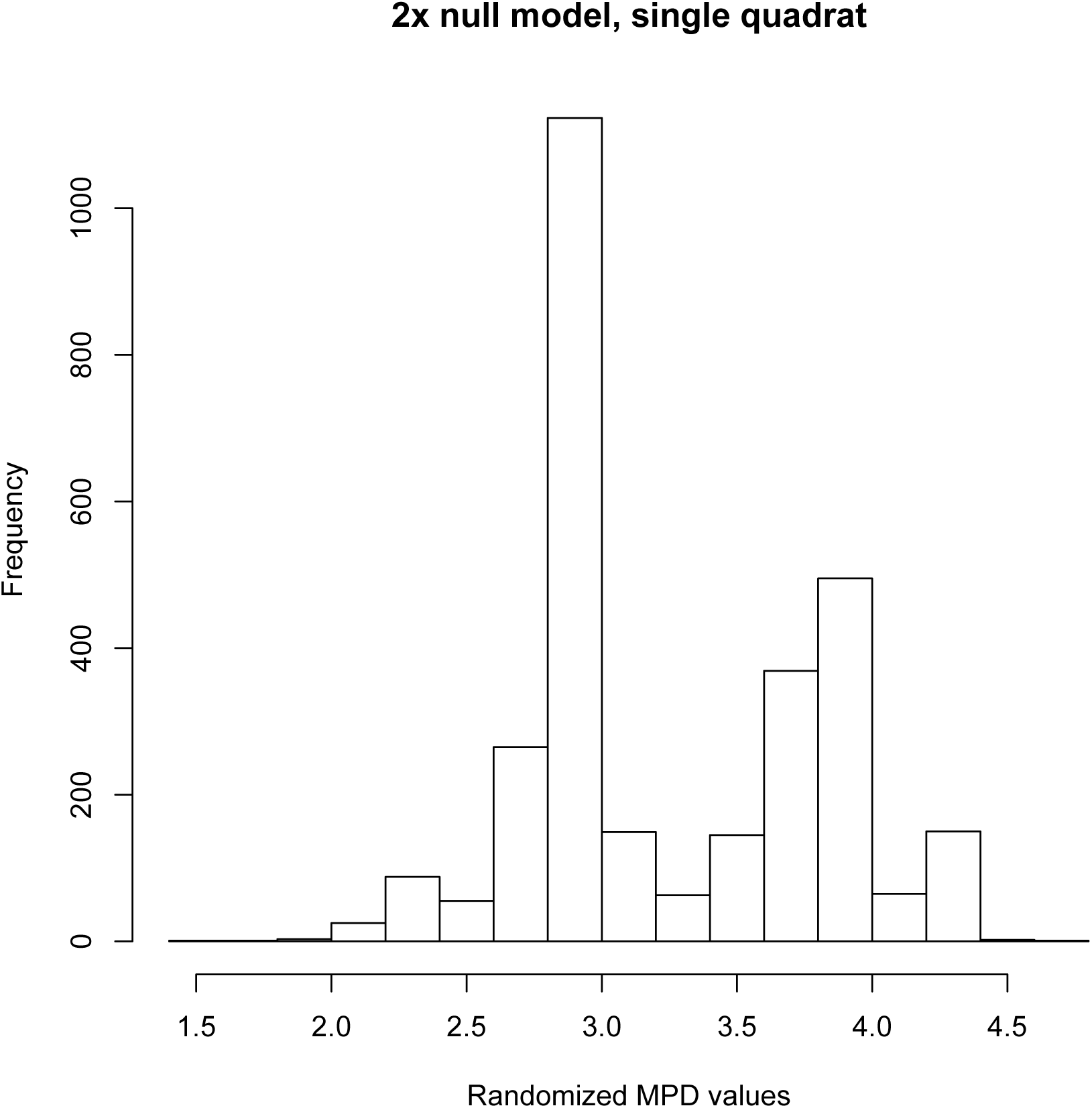
Distribution of randomized MPD values after 3,000 randomizations of the CDM described in this appendix. Here, all randomized values from one of the three quadrats that contained five species are shown (i.e. there are 3,000 randomized MPD values shown in the histogram).

**Figure S6.4.**
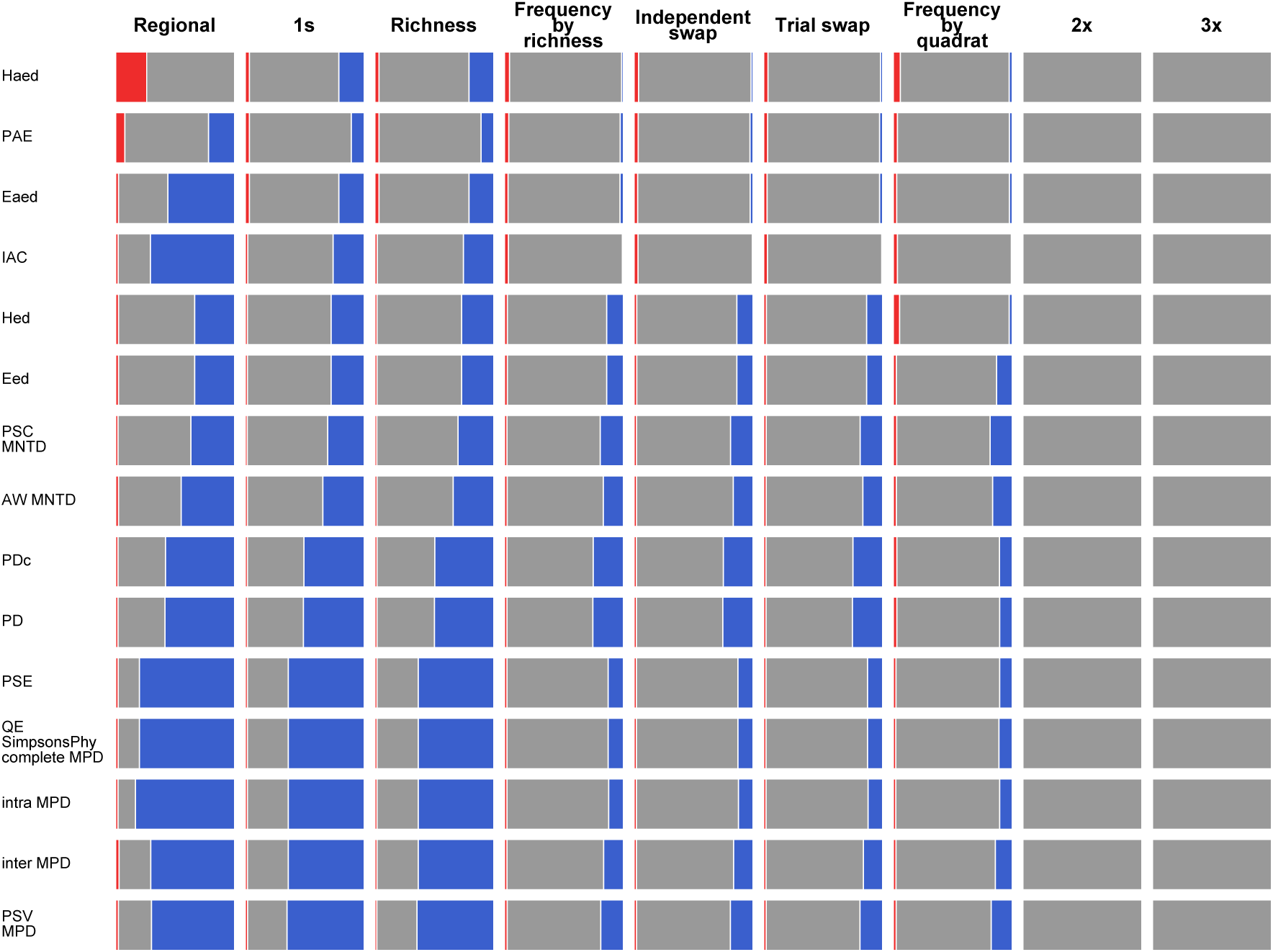
Overall performance of metric + null model approaches when randomizations were concatenated by quadrat and significance assessed on a per-quadrat basis. Red and gray bars summarize type I and II errors, respectively (see text for definition). Blue bars provide an indication of success. Metrics and nulls are arranged in order from overall best-performing to worst, with the best approaches in the bottom left corner.

**Figure S6.5.**
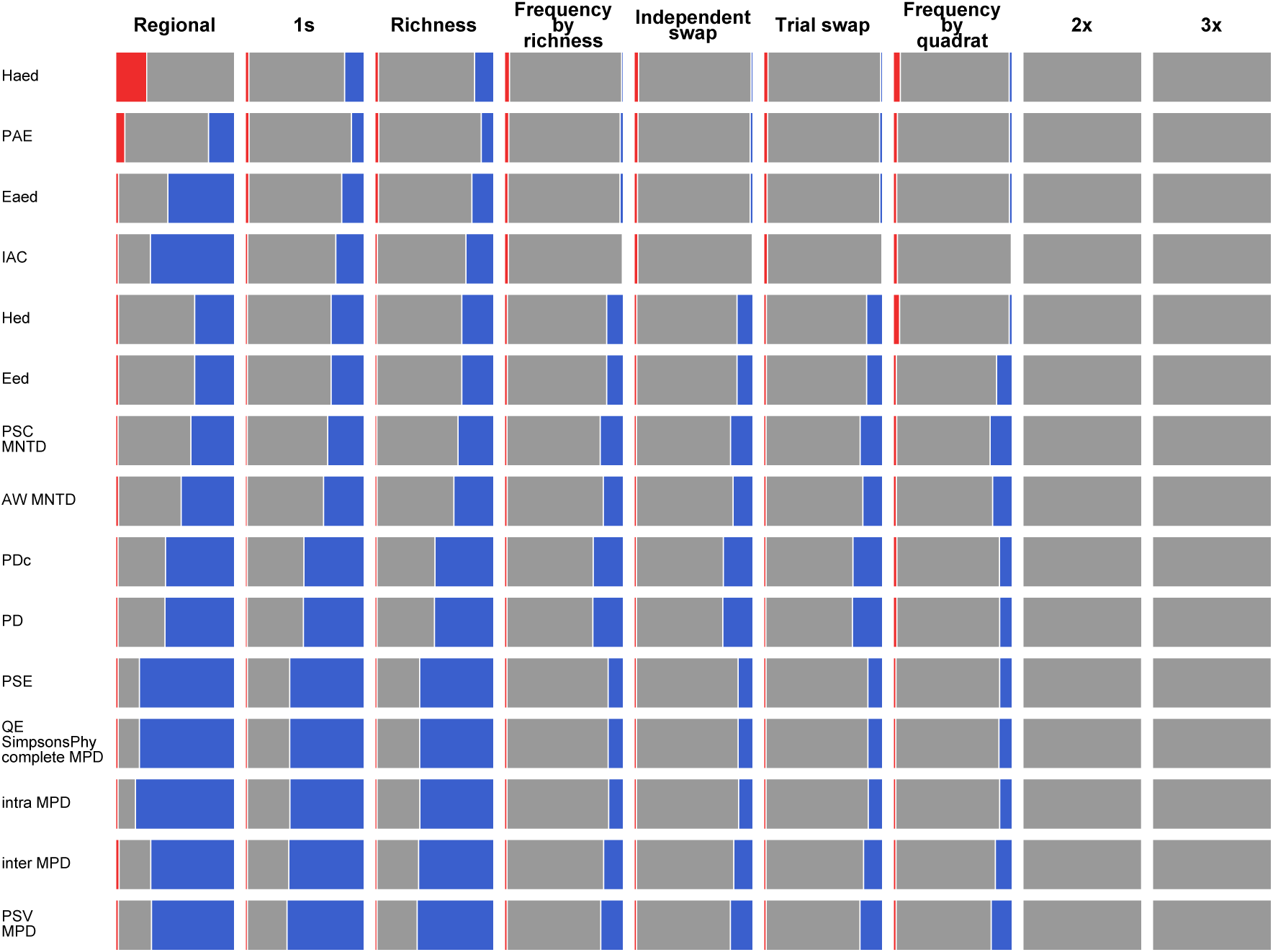
Overall performance of metric + null model approaches when randomizations were concatenated by richness (except for the frequency by quadrat model, which is always concatenated by quadrat) and significance assessed on a per-quadrat basis. Red and gray bars summarize type I and II errors, respectively (see text for definition). Blue bars provide an indication of success. Metrics and nulls are arranged in order from overall best-performing to worst, with the best approaches in the bottom left corner.

